# A phenotype-based forward genetic screen identifies *Dnajb6* as a sick sinus syndrome gene

**DOI:** 10.1101/2022.01.25.477752

**Authors:** Yonghe Ding, Di Lang, Jianhua Yan, Haisong Bu, Hongsong Li, Kunli Jiao, Jingchun Yang, Tai Le, Karl J. Clark, Stephen C. Ekker, Hung Cao, Yuji Zhang, Yigang Li, Alexey V. Glukhov, Xiaolei Xu

**Affiliations:** Department of Biochemistry and Molecular Biology, Department of Cardiovascular Medicine, Mayo Clinic, Rochester, MN, USA; The Affiliated Hospital of Qingdao University & The Biomedical Sciences Institute of Qingdao University (Qingdao Branch of SJTU Bio-X Institutes), Qingdao University, Qingdao, China; Department of Medicine, School of Medicine and Public Health, University of Wisconsin-Madison, Madison, WI, USA; Division of Cardiology, Xinhua Hospital Affiliated to Shanghai Jiaotong University School Of Medicine, Shanghai, China; Department of Cardiothoracic Surgery, Xiangya Hospital, Central South University, Changsha, China; Department of Cardiovascular Medicine, Jiading District Central Hospital Affiliated Shanghai University of Medicine & Health Science, Shanghai, China; Department of Electrical Engineering and Computer Science, University of California Irvine, Irvine, CA, USA; Department of Biochemistry and Molecular Biology, Mayo Clinic, Rochester, MN, USA; Department of Biomedical Engineering, University of California Irvine, Irvine, CA, USA; Department of Epidemiology and Public Health, University of Maryland School of Medicine, Baltimore, Maryland, USA

**Keywords:** Sinus arrest, Sick sinus syndrome, Dnajb6, Electrocardiogram, Zebrafish

## Abstract

Sick sinus syndrome (SSS) is a group of heart rhythm disorders caused by malfunction of the sinus node, the heart’s primary pacemaker. Partially owing to its aging-associated phenotypic manifestation and low expressivity, molecular mechanisms of SSS remain difficult to decipher. Here, we aim to develop a phenotype-based forward genetic approach in the zebrafish (*Danio rerio*) animal model for discovering essential genes which dysfunction could result in SSS-like phenotypes. Previously we showed the generation of protein trap library by using a revertible gene-breaking transposon (GBT)-based insertional mutagenesis system. Here, we reported the generation of a collection of 35 zebrafish insertional cardiac lines derived from this protein trap library, which was screened using electrocardiographic measurements. As a result, three mutants with SSS-like phenotypes were identified. We then focused on one of these 3 GBT mutants called *GBT411* in which *dnajb6b* gene was disrupted, and conducted expressional, genetic, transcriptome, and electrophysiological studies using both zebrafish and mouse models. These studies confirmed the identity of *Dnajb6* as a novel SSS causative gene with a unique expression pattern within the specialized population of sinus node pacemaker cardiomyocytes that lack the expression of HCN4 channels. Together, this study demonstrates the feasibility of a genetic screening approach in an adult vertebrate animal model for discovering new genetic factors for a heart rhythm disorder such as SSS.

## 1. Introduction

Cardiac arrhythmia affects >2% of individuals in community-dwelling adults.^1^ Sick sinus syndrome (SSS), also known as sinus node dysfunction or sinoatrial node (SAN) disease, is a group of heart rhythm disorders affecting cardiac impulse formation and/or propagation from the SAN, the heart’s primary pacemaker. SSS manifests a spectrum of presentations such as sinus pause or arrest (SA), bradycardia, sinoatrial exit block, or tachy-brady syndrome accompanied by atrial fibrillation (AF).^2, 3^ In addition, 20% to 60% SSS patients show abnormal response to autonomic stresses.^4^ SSS occurs most commonly in elderly, with an estimated prevalence of 1 case per 600 adults over age 65. Symptomatic SSS can lead to inadequate blood supply to the heart and body and contribute significantly to life-threatening problems such as heart failure and cardiac arrest. While SSS is the most common indication for pacemaker implantation worldwide,^5^ the mechanisms of SSS remain poorly understood, making it difficult to stratify SSS risk in vulnerable cohorts of patients and development of effective pharmacologic therapy for pacemaker abnormalities.

To develop mechanism-based diagnostic and therapeutic strategies for SSS, it is desirable to discover genes that are expressed in the SAN and may contribute to SSS. Unfortunately, very limited number of SSS genes and related animal models are currently available. While mutations in the cardiac sodium channel α-subunit encoding gene (*SCN5A*) ^6, 7^ and hyperpolarization-activated cyclic nucleotide-aged channel encoding gene (*HCN4*) ^8, 9^ have been found to cause SSS, only a few other genes affecting the structure and/or function of the SAN were identified to increase the risk of developing SSS.^10, 11^ Classic human genetic linkage analysis-based approach has played important roles in gene discovery, but it is largely limited by the availability of suitable pedigree, especially in this age-dependent disease.^12^ More recently, the genome-wide association studies (GWASs) have been used to identify novel genetic susceptibility factors associated with SSS.^11, 13^ However, owing to its statistic and associative nature, it has been difficult to confidently establish genotype-phenotype relationships for the vast amount of variants.^14, 15^ Alternative approaches for effective identification of essential genes for SSS are thus needed.

Phenotype-based forward genetic screen in model organisms is a powerful strategy for deciphering genetic basis of a biological process. Without any *a prior* assumption, new genes can be identified that shed light on key signaling pathways. However, this approach is difficult to carry out in adult vertebrates, because of significantly increased burden of colony management efforts.^16, 17^ To address this bottleneck, zebrafish, a vertebrate with higher throughput than rodents, has been explored to study cardiac diseases.^18^ Despite its small body size, a zebrafish heart has conserved myocardium, endocardium, and epicardium as found in human, and adult zebrafish shows strikingly similar cardiac physiology to humans.^19^ Its heart rate is around 100 beats per minute (bpm), which is much comparable to that in human than in rodents. Adult zebrafish models for human cardiac diseases such as cardiomyopathies have been successfully generated.^20^ Besides *N*-ethyl-*N*-nitrosourea (ENU)-based mutagenesis screens that have been conducted to identify embryonic recessive mutants, insertional mutagens such as those based on viruses and/or transposons have been developed to further increase the throughput of the screen, opening doors to screening genes affecting adult phenotypes.^21, 22^ Our team recently reported a gene-breaking transposon (GBT)-based gene-trap system in zebrafish which enables to disrupt gene function reversibly at high efficiency (>99% at the RNA level).^23^ Approximately 1,200 GBT lines have been generated, laying a foundation for adult phenotype-based forward genetic screens.^24^ Because the expression pattern of the affected genes in each GBT line is reported by a fluorescence reporter, we enriched GBT lines with cardiac expression and generated a zebrafish insertional cardiac (ZIC) mutant collection.^25^ Through stressing the ZIC collection with doxorubicin, an anti-cancer drug, we demonstrated that novel genetic factors of doxorubicin-induced cardiomyopathy (DIC), such as Dnaj (Hsp40) homology, subfamily B, member 6b (*dnajb6b*), sorbin and SH3 domain-containing 2b (*sorbs2b*) and retinoid x receptor alpha a (*rxraa*), could be successfully identified.^26-28^ Follow up studies on these hits confirmed their identity as important cardiomyopathy genes.

Encouraged by our success in identifying new genetic factors for DIC, we reasoned that genes for rhythm disorders could be similarly identified by directly screening adult ZIC lines using echocardiographic measurement. We had recently optimized a commercially available ECG system to define SA episodes in an adult zebrafish, and the baseline frequency of aging-associated SSS in wild-type (WT) adult zebrafish.^29^ Here, we reported a pilot screen of our ZIC collection using this ECG platform and the resultant discovery of 3 positive hits, followed by comprehensive expressional and functional analysis of *dnajb6b* gene that is linked to one of the hits. Together, our data prove the feasibility of a phenotype-based screening strategy in adult zebrafish for discovering new rhythm genes.

## 2. Results

### 2.1 Identification of 35 zebrafish insertional cardiac (ZIC) mutants

We recently reported the generation of more than 1,200 zebrafish mutant strains using the gene-break transposon (GBT) vector.^24^ The tagged gene in each GBT mutant is typically disrupted with 99% knockdown efficiency and its expression pattern is reported by a monomeric red fluorescent protein (mRFP) reporter.^24^ We screened 609 GBT lines based on their mRFP expression and identified 44 mutants with either the embryonic or adult heart expression.^26^ Then, we outcrossed these 44 lines, aided by Southern blotting to identify offsprings with a lower copy number of insertions,^25^ and identified 35 mutants with a single copy of the GBT insertion after 2-4 generations of outcross (Table 1).^26^ Using a combination of inverse PCR and/or 5’- and 3’-RACE PCR cloning approaches, we mapped the genetic loci of GBT inserts in these 35 mutants (Table 1).^25^ A majority of the affected genes have human orthologs with a corresponding Online Mendelian Inheritance in Man (OMIM) number. Because each GBT line contains a single GBT insertion that traps a gene with cardiac expression, these 35 GBT lines were termed as zebrafish insertional cardiac (ZIC) mutants.

**Table 1.**
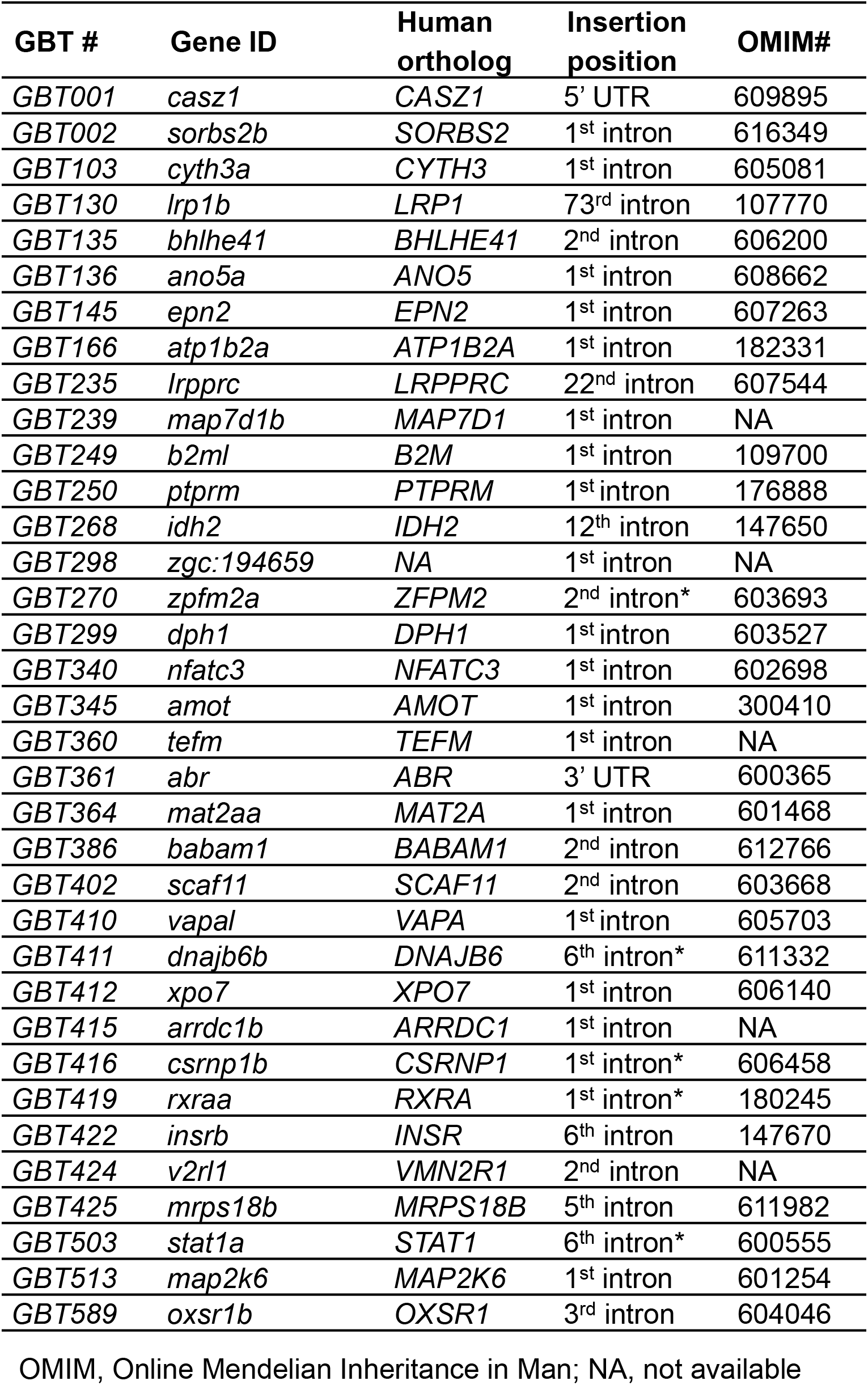
Collection of 35 zebrafish insertional cardiac (ZIC) mutants

### 2.2 An ECG screen of 35 ZIC lines identified 3 mutants with increased incidence of SA and/or AV block episodes

Because each ZIC mutant disrupts a gene with cardiac expression, we enquired whether an ECG screening can be conducted to identify genetic lesions that result in arrhythmia. Since aging is a strong risk factor for heart rhythm disorders, we carried our screen in fish aged from 1.5 to 2 years old to facilitate the manifestation of cardiac rhythm abnormalities. Because these fish are offsprings of incrosses and have been preselected based on the RFP tag, their genotype consists of both heterozygous and homozygous for the affected genes. In WT fish aged around 2 years old, we noted baseline SA episodes in about 1 out of 20 fish (5%) fish.^29^ By contrast, among the 35 ZIC lines, we noted a significantly increased incidence of SA in 3 lines, including 3 out of 13 *GBT103* fish at 1.5 years old, 4 out of 10 *GBT410* fish at 2 years old, and 3 out of 8 *GBT411* fish at 2 years old (Figure 1A). In addition to SA, we also noted incidence of atrioventricular block (AVB) in 4 out of 13 *GBT103* fish at 1.5 years of age. Because the increased incidence of SA and/or AVB is hallmark of SSS, these 3 lines were thus identified as 3 candidate SSS-like mutants.

**Figure 1.**
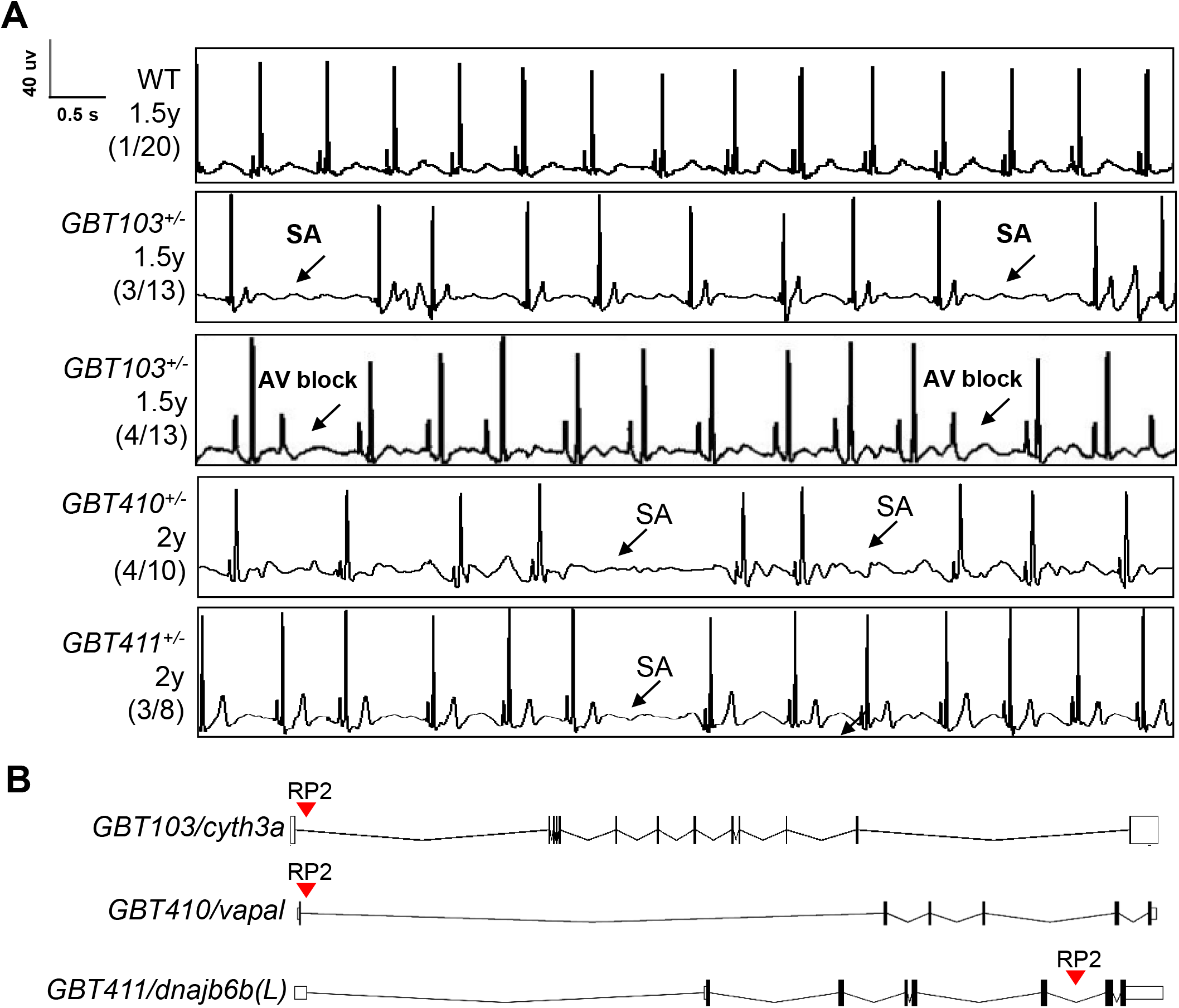
Screening of 35 ZIC lines identified 3 mutants with increased incidence of SA and/or AVB episodes. **(A)** Representative ECG recordings for 3 heterozygous GBT mutants with incidence of sinus arrest (SA) and/or atrioventricular block (AVB) episodes compared to WT control. **(B)** RP2 genebreak transposon insertional positions in the 3 candidate SSS mutants.

To confirm the linkage between genetic lesions and the SSS-like phenotypes, we incrossed these 3 ZIC mutants to obtain homozygous animals. This is possible because the precise insertional positions for all the 35 ZIC lines have been mapped (Table 1, Figure 1B). We carried genotyping PCR to identify homozygous mutants for the 3 candidate ZIC lines using genomic DNA isolated from their tail fins, raised up homozygous fish to 16 months, and carried out ECG assays at room temperature (25 °C). In contrast to 5% WT fish whereby SA episodes can be detected, significantly increased SA incidence was noted in all 3 homozygous mutants, with an incidence of 57.1% in the *GBT103/cyth3a*, 44.4% in the *GBT410/vapal*, and 40% in the *GBT411/dnajb6b* mutant, respectively (Table 2). We also noted a reduced heart rate, another SSS phenotypic trait in the *GBT411*^*-/-*^ homozygous, but not the other two GBT homozygous mutants (Table 2).

**Table 2.** ECG quantification to validate 3 GBT lines as SA mutants in homozygous fish

| Genotype | Age | N | SA incidence (%) | SA Frequency (epm) | Heart rate (bpm) |
| --- | --- | --- | --- | --- | --- |
| WT | 16 m | 20 | 5.0 | 2.0 | 100.1±11.1 |
| *GBT103<sup>-/-</sup>* | 16 m | 7 | 57.1* | 3.4±2.9 | 89.1±18.8 |
| *GBT410<sup>-/-</sup>* | 16 m | 9 | 44.4* | 1.4±0.6 | 99.9±17.7 |
| *GBT411<sup>-/-</sup>* | 16 m | 10 | 40.0* | 3.0±1.0 | 90.6±7.5* |

SA, sinus arrest. bpm, beats per minute. N=7-20. \*, $P<0.05$ , data are expressed as mean±SEM. For SA incidence comparison, Chi-square test. For heart rate comparison, unpaired student's *t*-test.

To seek additional evidence supporting our screening strategy, we decided to focus on the *GBT411/dnajb6b* mutant that is the most arrhythmogenic – this mutant is also characterized with reduced heart rate phenotype. Because arrhythmic mutants often manifest an aberrant response to extrinsic regulation of the heart rate, we examined responses of the *GBT411/dnajb6b* homozygous mutants (*GBT411*^*-/-*^) to autonomic stimuli by stressing them with 3 compounds, including isoproterenol, a β-adrenoreceptor agonist for sympathetic nervous system; atropine, an anticholinergic inhibitor; and carbachol, a cholinergic agonist for parasympathetic nervous system. After administrating these drugs to the *GBT411*^*-/-*^ fish at 1 year old via intraperitoneal (IP) injection, we noted aberrant heart rate response to both atropine and carbachol, while its response to isoproterenol remained unchanged (Supplemental Figure 1). Next, we stressed the *GBT411*^*-/-*^ fish with verapamil, an L-type Ca^2+^ channel antagonists, to stress out cardiac pacemaking and unmask SSS phenotype. Indeed, SA incidence was significantly increased in the *GBT411*^*-/-*^ fish at 10 months of age (Supplemental Table 1). Similarly, the heart rate was significantly reduced in the *GBT411*^*-/-*^ fish compared to WT controls. Together, these data provided additional evidence to support *GBT411/dnajb6b* as an arrhythmia mutant.

### 2.3 Dnajb6 expression is enriched in the SAN tissue, manifesting a unique expression pattern

*Dnajb6* was previously identified as a cardiomyopathy-associated gene,^26^ raising concerns on whether the arrhythmic phenotype in the *GBT411/dnajb6b* mutant is a primary defect in the cardiac conduction system or a consequence of cardiac remodeling in cardiomyocytes. To address this further, we firstly defined the expression of the Dnajb6 protein in the heart. Our previous characterization of the mRFP reporter in the *GBT411/dnajb6b* fish revealed expression of Dnajb6b protein in both the embryonic and the adult hearts.^25, 26^ To enquire its expression in the cardiac conduction system (CCS), we crossed the *GBT411/dnajb6b* line into the sqET33-mi59B transgenic line in which EGFP labels the zebrafish SAN and atrio-ventricular canal (AVC) cells.^30^ Co-localization analysis demonstrated that the mRFP positive, Dnajb6b-expressing cells partially overlap with the EGFP signal labeling SAN cells at the base of atrium in the embryonic heart at 3 days post-fertilization and also AVC in adult heart tissues (Figure 2A and 2B). It should be noted that the Dnajb6b-mRFP-positive expression patterns overlap with but extend beyond the sqET33-mi59B EGFP-positive expression patterns in both embryonic and adult fish hearts (Figure 2A and 2B).

**Figure 2.**
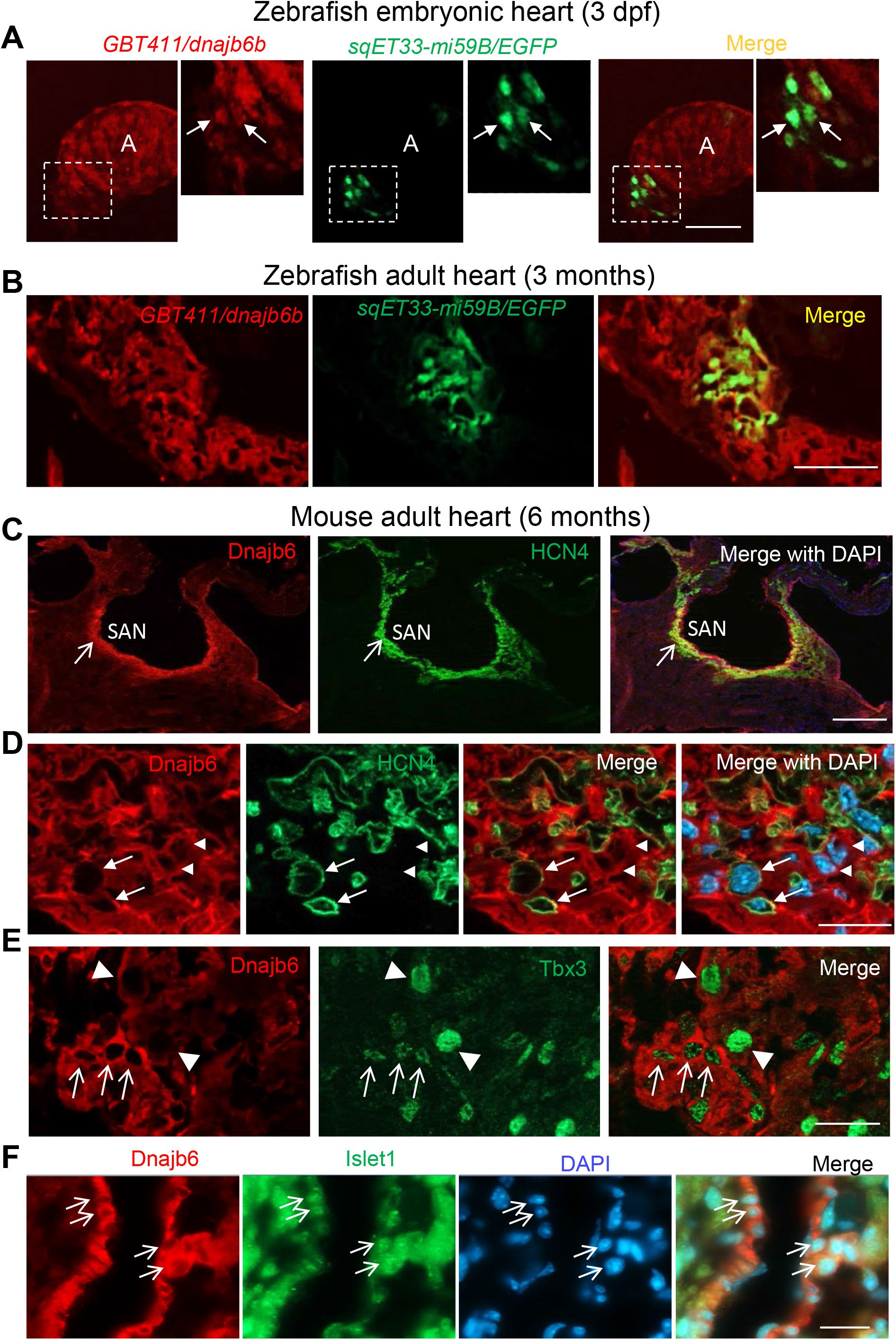
Dnajb6 is expressed in the SAN in both zebrafish and mouse. **(A-B**) Co-localization analysis of mRFP in GBT411/dnajb6b with the reporter line sqET33-mi59B in which EGFP labels cardiac conduction system (CCS) in zebrafish. The mRFP reporter for the GBT411 tagged Dnajb6b protein partially overlaps with the EGFP reporter in the sqET33-mi59B transgenic line that labels the SAN in embryonic atrium at 3 dpf **(A)** and atrio-ventricular canal (AVC) in adult hearts **(B)**. Arrows indicate EGFP+ cells in the sqET33-mi59B reporter line. A: atrium. V: ventricle. dpf, days post-fertilization. **(C)** The Dnajb6 antibody immunostaining signal largely overlapped with the HCN4 immunostaining signal in mouse SAN tissues under low magnification. **(D)** Under higher magnification, expression of Dnajb6 (red) only partially overlapped with HCN4 (green) as revealed by antibody co-immunostaining. Arrows point to cells with overlapping patterns. Arrowheads point to cells with no-overlapping. **(E)** Shown are images indicating expression of Dnajb6 protein largely overlapping with the Tbx3 antibody immunostaining with medium to low intensity, but not strong signal in the SAN tissues. Arrows point to cells with strong Tbx3 immunostaining signal. Arrowheads point to cells with medium to low level of Tbx3 immunostaining signal. **(F)** Shown are images indicating expression of Dnajb6 protein largely overlapping with the Islet1, a transcription factor labeling SAN cells. Arrows point to cells with overlapping immunostaining signal for both Dnajb6 and Islet1. Scale bars in A, C, 50 μm. Scale bars in B, D, E, F, 20 μm.

To seek additional evidence supporting expression and function of Dnajb6 in the CCS, we turned to the mouse model, and noted Dnajb6 protein expression in all 4 cardiac chambers in a sectioned mouse heart tissue (Supplemental Figure 2). Interestingly, we found a highly enriched expression of Dnajb6 specifically in the SAN region, as indicated by its localization in the region with expression of HCN4 channel which are responsible for the generation of hyperpolarization-activated pacemaker “funny” current in pacemaker cells (Figure 2C). However, at higher magnification images, only a small proportion of Dnajb6-positive cells showed colocalization with the HCN4-positive cells (arrows for colocalized cells vs. arrowheads non-colocalized cells in Figure 2D). In addition, we noted co-localization of Dnajb6 with Tbx3 and Islet1, two transcription factors that specify the formation of the SAN cells.^32^ Interestingly, we found a negative correlation between Dnajb6 and Tbx3 expression levels: cells with strong Dnajb6 expression tend to overlap with cells that show weak Tbx3 signal, while cells with weak Dnajb6 expression tend to overlap with the cells with strong Tbx3 signal. In contrast, the Dnajb6 expressing cells largely overlap with the Islet1 positive cells. In summary, the enriched expression of Dnajb6 in the SAN region may indicate that Dnajb6 could contribute to SSS development; however, its unique expression patterns underscored heterogeneity of pacemaker cells within the SAN.^33, 34^

### 2.4 The *Dnajb6*^*+/-*^ mice manifest features of SSS when there is no sign of cardiomyopathy

To test the conservation of the cardiac arrhythmic functions of dnajb6 suggested from zebrafish, we obtained a global *Dnajb6* knock out (KO) mouse line. The mutant harbors a deletion of 36,843 bp nucleotides spanning from the first intron to the last intron of *Dnajb6* gene located in the Chromosome 5, which was created by the insertion of the Velocigene ZEN-Ub1 cassette and subsequent LoxP excision using Cre (Figure 3A). Genotyping PCR using a combination of the *Dnajb6* gene-specific and the Zen-Ubi cassette-specific primers was carried out to identify both Dnajb6 heterozygous (*Dnajb6*^*+/-*^) and homozygous (*Dnajb6*^*-/-*^) KO mice (Figure 3B). At the protein level, both the Dnajb6 short (S) and long (L) isoforms were reduced by ∼ 50% in *Dnajb6*^*+/-*^ mouse hearts (Figure 3C), and near completely depleted in *Dnajb6*^*-/-*^ mutant hearts. Consistent with a previous report,^35^ *Dnajb6*^*-/-*^ KO mice were embryonic lethal, likely due to the placental defects (data not shown). The *Dnajb6*^*+/-*^ mice were able to grow to adulthood without visually noticeable phenotypes until at least 1 year of age. Cardiac mechanical function remained normal, as indicated by indistinguishable cardiac echocardiography indices from those of WT siblings at the same age (Table 3). However, significantly increased frequency of SA and AVB episodes, as well as bradycardia phenotype, were noted in the *Dnajb6*^*+/-*^ mice at 6 months old (Figure 3D and 3E, and Table 4). Similar to the *GBT411/dnajb6b* mutant in zebrafish, *Dnajb6*^*+/-*^ mice exhibited an impaired response to autonomic stimuli including isoproterenol and carbachol (Figure 3E). Together, these studies suggest that *Dnajb6*^*+/-*^ mice manifest SSS phenotype without structural/functional remodeling of the heart.

**Figure 3.**
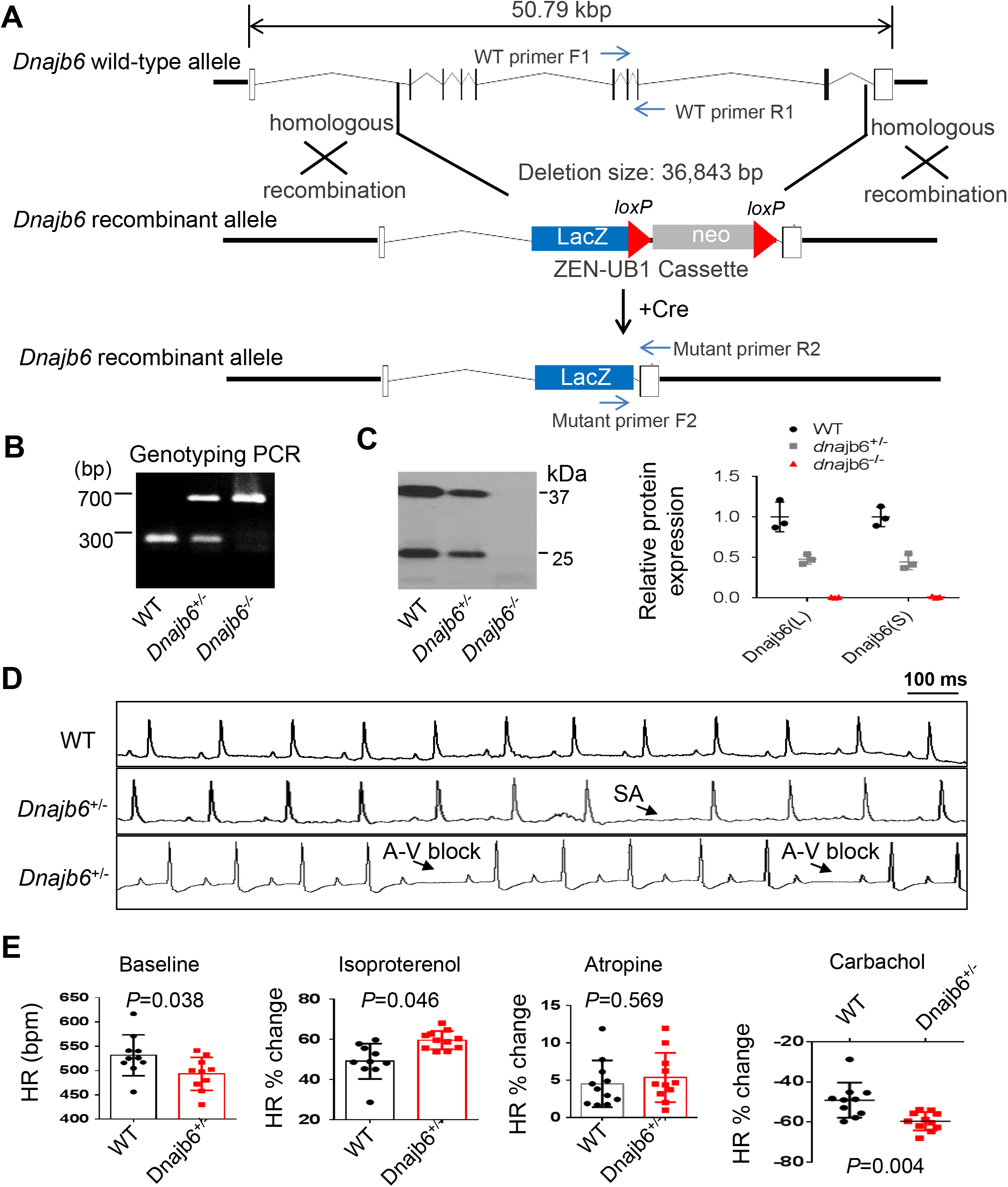
*Dnajb6*^*+/-*^ mice exhibited increased incidence of SA and AVB and impaired response to autonomic stimuli. **(A)** Schematics of the Dnajb6 knockout (KO) mice. The insertion of Velocigene cassette ZEN-Ub1 created a deletion of 36,843 bp nucleotides spanning from the first to the last intron of the Dnajb6 gene at the Chromosome 5. The neomycin selection cassette was excised after crossed to a Cre expression line. **(B)** Representative DNA gel images of PCR genotyping for identifying WT (300 bp), *Dnajb6*^*+/-*^ heterozygous (hets), and Dnajb6-/- homozygous (homo) mutant alleles (700 bp). **(C)** Western blotting and quantification of Dnajb6 short (S) and long (L) protein expression in WT and Dnajb6 mutants. N=3 animal per group. **(D**) Shown are representative ECG recordings results showing SA and AVB phenotypes detected in the *Dnajb6*^*+/-*^ mice at 6 months. **(E)** The *Dnajb6*^*+/-*^ mice manifests impaired response to different autonomic stimuli. N=10-12 mice per group. Unpaired student’s t-test. SA, sinus arrest. AVB, atrioventricular block.

**Table 3.**
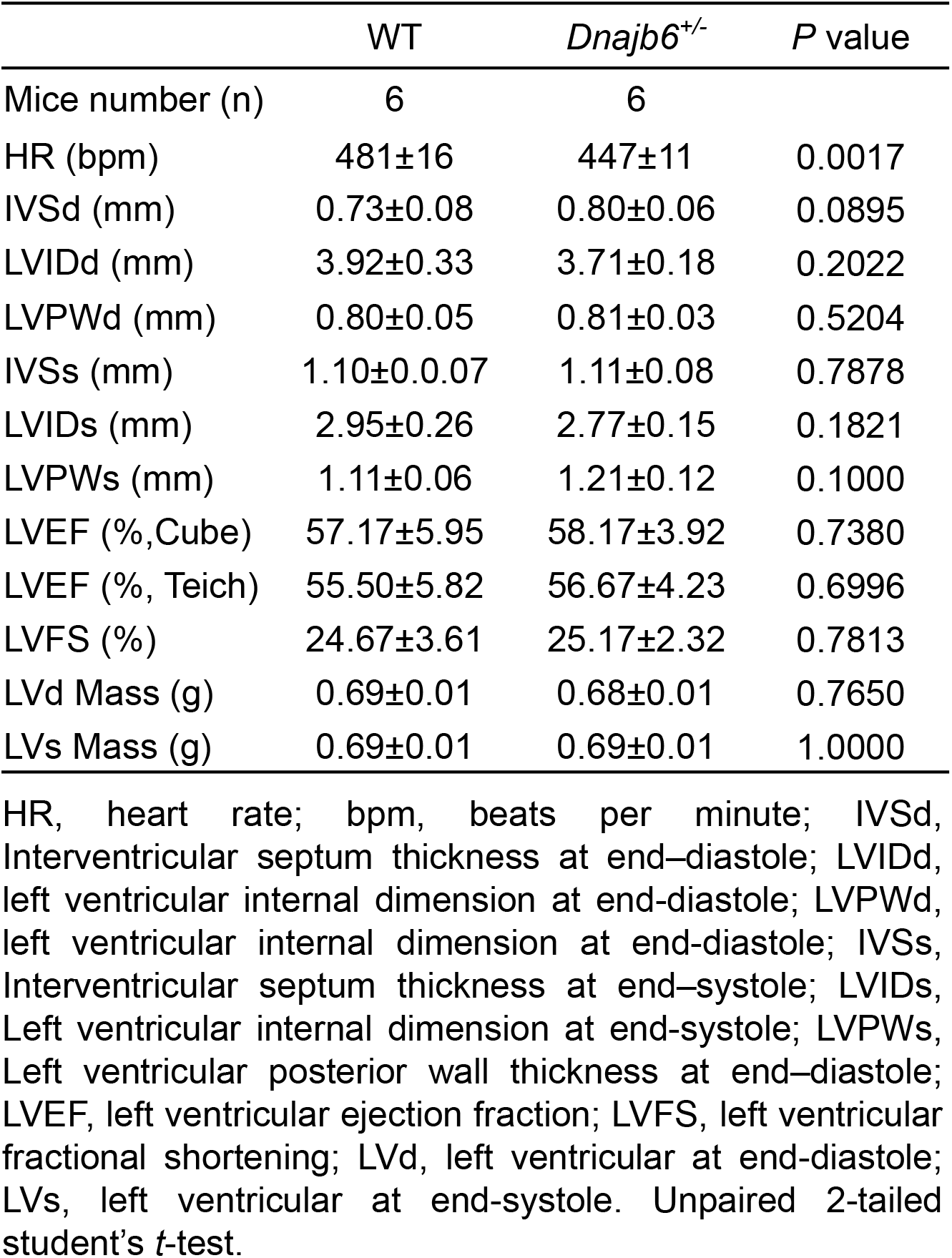
Echocardiography indices in the *Dnajb6*^*+/-*^ heterozygous mice compared to WT controls at 1 year

**Table 4.**
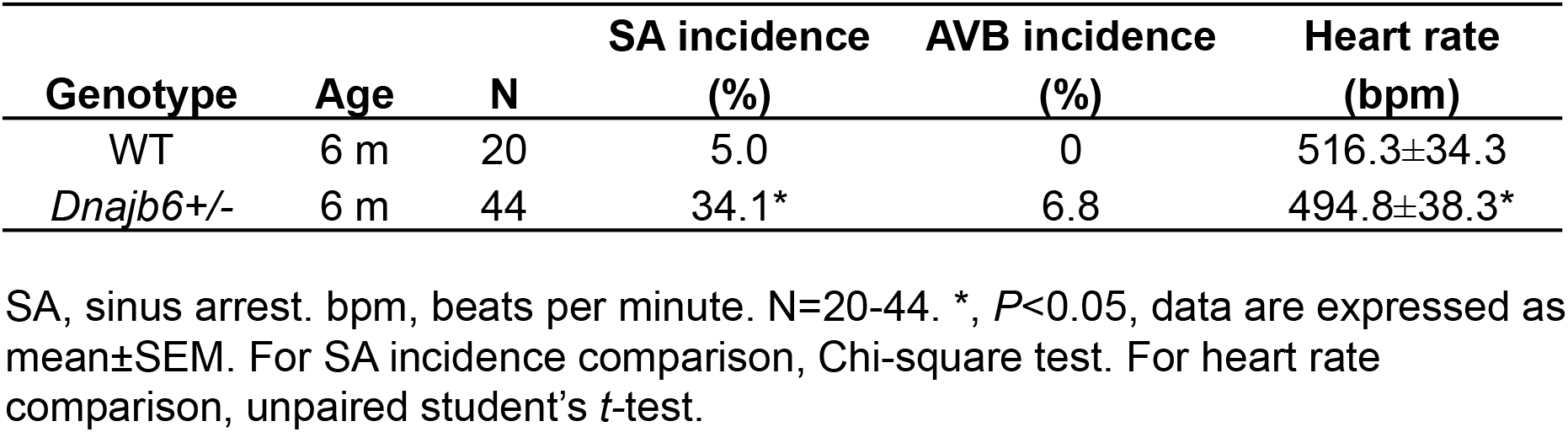
ECG quantification of *Dnajb6* heterozygous mice at 6 months of age

### 2.5 *Ex vivo* evidences of SAN dysfunction in the *Dnajb6*^*+/-*^ mice

To prove SAN dysfunction in *Dnajb6*^*+/-*^ mice, we performed electrophysiological assessment of SAN pacemaker function by high-resolution fluorescent optical mapping of action potentials from isolated mouse atria at 1 year of age. We firstly analyzed the distribution of the leading pacemaker location site in *Dnajb6*^*+/-*^ mice compared to WT control. In WT mice, leading pacemakers were mostly located within the anatomically and functionally defined SAN region (Figure 4A and 4B).^18, 36-38^ In contrast, significant increase in the number of leading pacemakers located outside of the SAN, including the subsidiary atrial pacemakers and inter-atrial septum pacemakers, was observed in *Dnajb6*^*+/-*^ mice. In addition, in *Dnajb6*^*+/-*^ mice, we also found a highly irregular heart rate, accompanied by the presence of multiple competing pacemakers and a beat-to-beat migration of the leading pacemaker between various sites which included SAN, right atrial ectopic (subsidiary) pacemakers, and inter-atrial septum (Figure 4C and 4D). Similar to the results from the *in vivo* studies, bradycardia phenotype was consistently detected in the isolated atrial preparations as well (Figure 4E). Optical mapping on isolated atrial preparations further revealed different responses of heart rate during isoproterenol, atropine, and carbachol stimulations in *Dnajb6*^*+/-*^ mice. Significantly increased cycle length (CL) variations were also observed at baseline and upon carbachol stimulation (Figure 4F).

**Figure 4.**
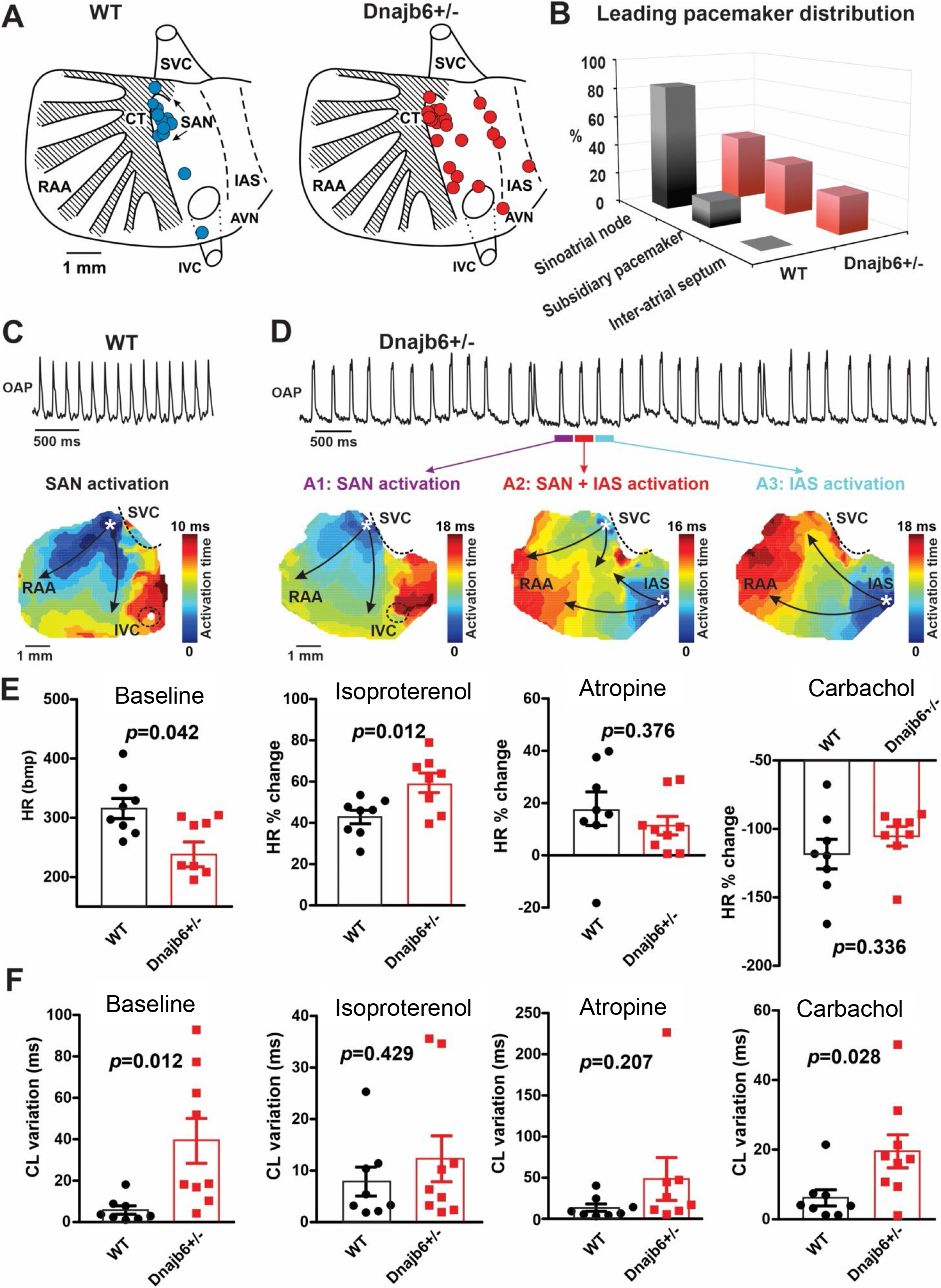
SAN dysfunction in the *Dnajb6*^*+/-*^ mice. **(A)** Leading pacemakers were located and plotted from both WT (blue dots) and *Dnajb6*^*+/-*^ (red dots) mice. One mouse could have multiple leading pacemaker locations due to the competing pacemakers and ectopic activities. SVC and IVC, superior and inferior vena cava; RAA, right atrial appendage; CT, crista terminalis; IAS, inter-atrial septum; AVN, atrioventricular node. Distribution of the leading pacemakers is summarized in panel. **(B)** Majority of leading pacemakers located within the SAN area in WT whereas, in *Dnajb6*^*+/-*^ mice, significant increase of leading pacemakers locating in subsidiary pacemaker area and IAS was observed. **(C-D)** Activation map based on the optical mapping of action potentials showed representative leading pacemaker locations in WT (SAN) and *Dnajb6*^*+/-*^ mice (SAN and IAS areas). **(E)** Optical mapping on isolated atrial preparation showed bradycardia (baseline) and different responses of heart rate during isoproterenol, atropine, and carbachol stimulations between WT and *Dnajb6*^*+/-*^ mice. N=7-9 mice per group. Unpaired student’s t-test. **(F)** Increased cycle length (CL) variation was observed in *Dnajb6*^*+/-*^ isolated atrial preparations during different autonomic stimulations. N=7-9 mice per group. Unpaired student’s *t*-test.

Importantly, in *Dnajb6*^*+/-*^ mice, we found significant prolongation of the SAN recovery time corrected to beating rate (cSANRT) measured both at baseline and under autonomic stresses, including stimulation by isoproterenol, carbachol, and atropine (Figure 5), confirming the presence of SAN dysfunction in *Dnajb6*^*+/-*^ mice. Optical mapping also showed that, unlike WT, the first spontaneous post-pacing atrial beats during SANRT measurements in *Dnajb6*^*+/-*^ mice were originated from ectopic locations outside of the SAN (Figure 5A-B), further supporting a suppressed SAN function.

**Figure 5.**
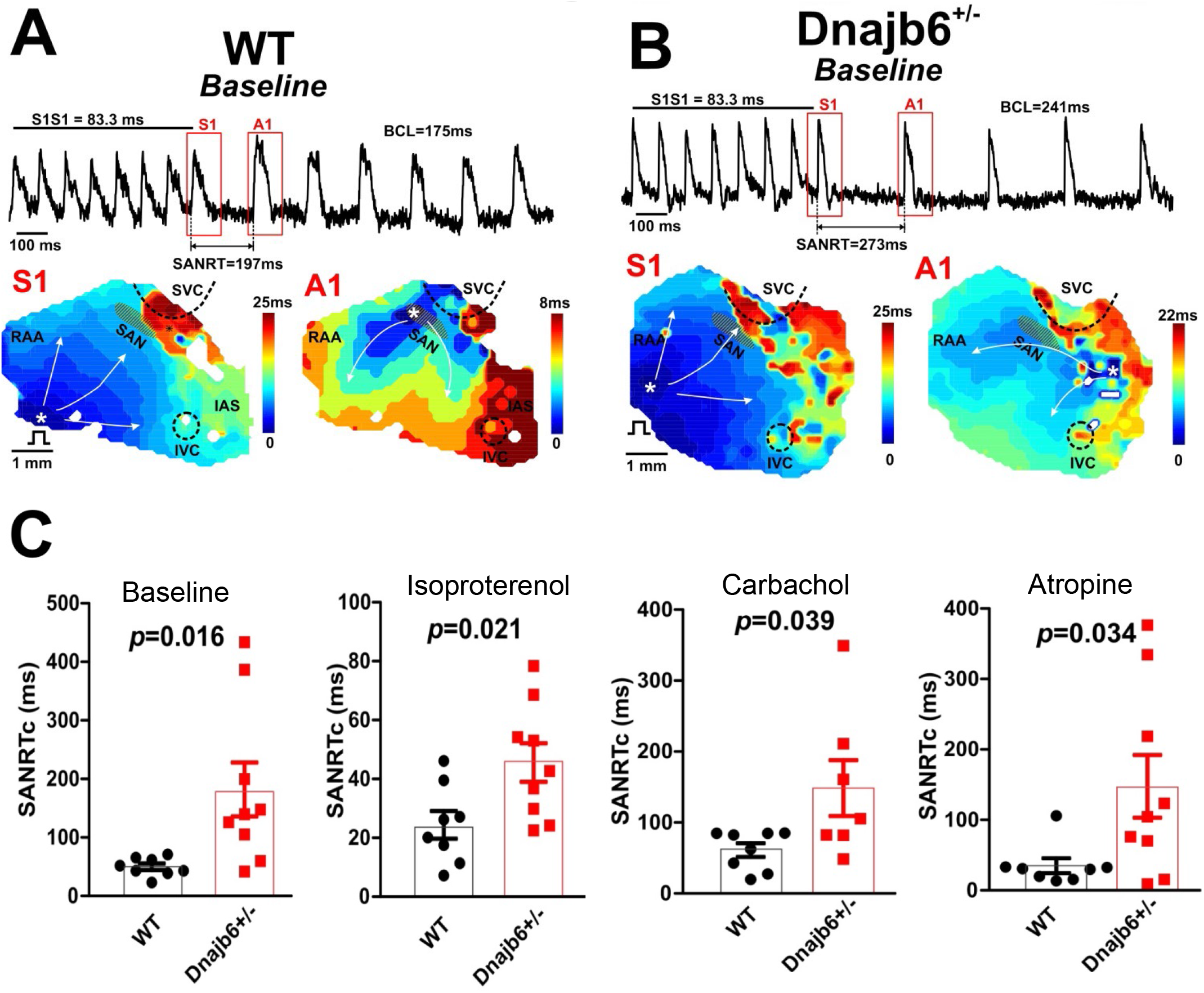
Sinus node recovery time was prolonged in the *Dnajb6*^*+/-*^ mice. **(A-B)** Representative activation maps reconstructed for the last pacing stimulus (S1) and the first spontaneous post-pacing atrial beat (A1) during SAN recovery time (SANRT) measurements are shown. A site of the earliest atrial activation is labeled by a white asterisk. In *Dnajb6*^*+/-*^ group, unlike WT, the first spontaneous post-pacing atrial beat (A1) originated from an ectopic location outside of the anatomically and functionally defined SAN area. **(C)** Summarized data for corrected SANRT (SANRTc) measured during different autonomic stimulations is shown. N=7-9 mice per group. Unpaired student’s *t*-test.

### 2.6 Transcriptome analysis of the *Dnajb6*^*+/-*^ mutant hearts identifies altered genes encoding ion channels and proteins in the Wnt/beta-catenin pathway

To seek molecular mechanisms underlying the SSS phenotypes observed in *Dnajb6*^*+/-*^ mice, we performed whole transcriptome RNA-sequencing experiments using right atrial tissues isolated from *Dnajb6*^*+/-*^ and WT mice at 1 year of age. Transcriptomes of biological replicates for *Dnajb6*^*+/-*^ mice did form a cluster that differs from the cluster for WT control samples, as indicated by principal component analysis (PCA) (Supplemental Figure 3A). Based on a cut-off of adjusted *P* value<0.05, 107 differentially expressed (DE) genes were identified, among which 37 genes were upregulated and 70 genes were downregulated in the *Dnajb6*^*+/-*^mutants compared with WT controls (Supplemental Figure 3B and 3C). Through Ingenuity pathway analysis (IPA), several diverse signaling pathways were identified to be altered in the *Dnajb6*^*+/-*^ mice (Supplemental Figure 3D). Among these 107 differentially expressed genes, we noted calcium handling related protein-encoding genes like *Slc24a2 and Cdh20*, ion channel-encoding genes including *Slc9a3r1, Kcnh7, Fxyd5 and Gjb5* (Figure 6A), as well as 4 Wnt pathway related genes (Figure 6B). We then performed quantitative RT-PCR analysis and experimentally confirmed dysregulation of these genes in the *Dnajb6*^*+/-*^ mice (Figure 6C). The data on calcium handling and ion channel-encoding genes are in line with the SAN dysfunction phenotype observed in the *Dnajb6*^*+/-*^ mice. Because Wnt signaling has been shown to direct pacemaker cell specification during SAN morphogenesis,^39, 40^ the identification of 4 Wnt pathway related genes could also support the observed SAN dysfunction phenotype in the *Dnajb6*^*+/-*^ mice.

**Figure 6.**
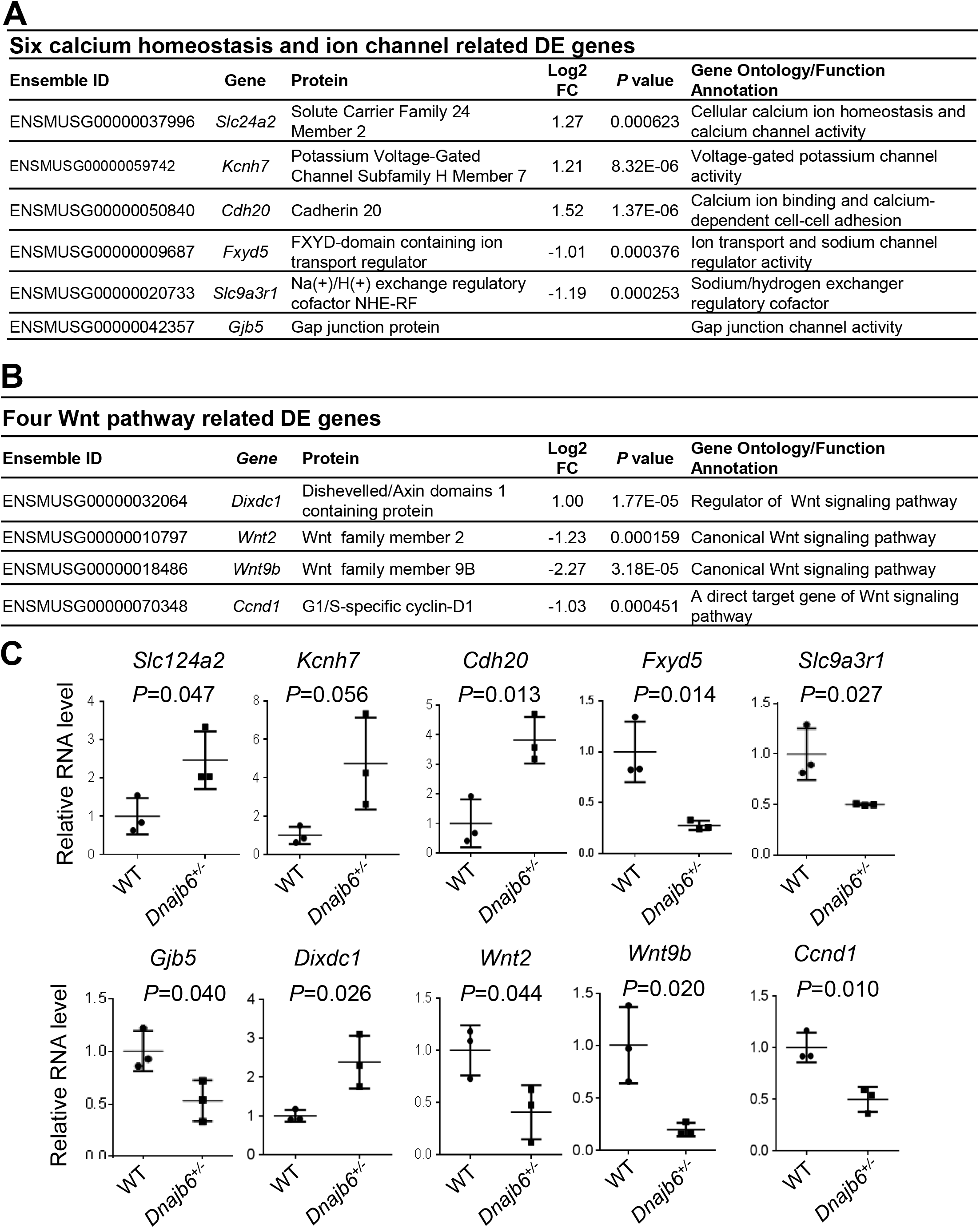
Transcriptomes are altered in the atrium of *Dnajb6*^*+/-*^ mice. **(A)** Expression of six calcium homeostasis and ion channel related genes were altered in the *Dnajb6*^*+/-*^ mice atrium. **(B)** Expression of four Wnt pathway related genes were altered in the *Dnajb6*^*+/-*^ mice atrium. **(C)** Quantitative polymerase chain reaction (qPCR) validation of DE genes listed in A and B, normalized to Gapdh; RNA was extracted from an individual moue atrium, which was considered a single biological replicate. Samples were collected in triplicate. N=3 mice per group. Unpaired student’s *t*-test.

## 3. Discussion

### 3.1 GBT lines enable a phenotype-based screening approach for discovering new SSS genes

This work is based on recent establishment of a GBT protein trap-based insertional mutagenesis screening strategy and the generation of a collection of 1,200 zebrafish mutant strains.^24^ Here, we demonstrated the feasibility of screening these GBT lines for discovering new genetic factors for SSS, an aging-associated human disease. To overcome the challenge of colony management efforts that is associated with an adult screen, we leveraged the following unique advantages of the GBT vectors and zebrafish models. First, the knockdown efficiency for the tagged gene in each GBT homozygous mutant is consistently high, which is typically >99%, which ensued the success of an adult screen. Second, because of a fluorescence tag, heterozygous GBT fish can be easily identified under a fluorescent microscope without the need of genotyping. As a consequence, a cardiac expression-based enrichment strategy can be used to identify ZIC lines. Instead of screening 609 GBT lines, only 35 ZIC lines need to be screened, which significantly reduced the workload. We acknowledge that some genes with extremely weak cardiac expression might be missed; however, this is not a concern during the early phase of a genome-wide screen. Third, it is economically feasible to house hundreds of mutant fish lines with different genetic lesions to 1-3 years old. Finally, we optimized an ECG technology, defined the baseline SSS in WT fish, and implemented heat-stress to zebrafish at old ages, which shall increase the SSS phenotypic expressivity.

While the forward genetic screening approach has been successfully utilized to pinpoint genetic basis of cardiogenesis in embryonic fish and doxorubicin-induced cardiomyopathy (DIC) in adult zebrafish,^21, 26^ this study extended this powerful genetic approach to adult zebrafish for discovering genetic factors associated with rhythm disorders. Given very little knowledge of molecular underpinnings of SSS, the development of this novel approach is significant. Human genetics approach has been difficult, partially owing to the aging associated nature - SSS-like phenotypes at its early stage are often missed, because SA episodes cannot be detected if the ECG measurement only covers a short time window. It takes years in patients to develop from asymptotic to onset of SSS symptoms. Moreover, human genetic studies of SSS are typically confounded by complicated environmental factors, which are minimalized in our zebrafish forward genetic approach - each ZIC mutant is maintained in a well-controlled living environment, and the only difference among different ZIC lines is a single genetic deficiency.

### 3.2 *Dnajb6* is a new SSS gene with a unique expression in SAN

The human *DNAJB6* gene encodes a molecular chaperone protein of the heat shock protein 40 (Hsp40) family. Dnajb6 has been previously linked to neurodegenerative diseases via its function in protein folding and the clearance of polyglutamine stretches (polyQ),^41, 42^ and to muscular dystrophy via its protein-protein interaction with Bag3 in the sarcomere.^43^ Our previous forward genetic screen in adult zebrafish identified *GBT411/dnajb6b* as a deleterious modifier for DIC.^26^ Here, we provided several evidences in both fish and mouse models, suggesting new functions of *Dnajb6* as a genetic factor for arrhythmia/SSS. First, *GBT411/dnajb6b* is one of three ZIC lines with SSS-like phenotypes that were identified from a screen of 607 GBT lines that is independent of the previous DIC screen. Second, in zebrafish, the increased incidence of SA episodes and reduced heart rate, two main features of SSS, were detected in 10 monthold *GBT411/dnajb6b* homozygous fish, when structural remodeling/cardiac dysfunction have not yet occurred.^25^ Similarly, bradycardia and SA episodes were noted in *Dnajb6*^*+/-*^ heterozygous KO mice at 6 months old, when the echocardiography indices remained indistinguishable from their age-matched siblings. Third, consistent with loss-of-function studies, Dnajb6 expression was detected in the SAN of both zebrafish and mice. Importantly, Dnajb6 is highly enriched in the SAN region of the mouse comparing to the surrounding atrial tissue. Fourth, transcriptome analysis of *Dnajb6*^*+/-*^ mice uncovered altered expression of genes involved in calcium handing, ion channels, and Wnt signaling pathway, which have been linked to the formation/function of the SAN. Together, we conclude that SSS might not be a consequence of Dnajb6 cardiomyopathy. Instead, the irregular heartbeat is most likely a direct consequence of Dnajb6 depletion in pacemaker cells, subsequently contributing to the pathogenesis of cardiomyopathy that occurs later. To ultimately confirm this hypothesis and to discern functions of Dnajb6 in SAN pacemakers from working cardiomyocytes, a tissue-specific KO line for Dnajb6 needs to be generated and studied.

Detailed examination of Dnajb6 expression in the SAN uncovered unique expression patterns. While Dnajb6 is highly expressed in the SAN and overlaps with one of the main SAN progenitors ISL1 (Figure 2C and 2F), we found a poor co-expression with one of the main pacemaker protein HCN4: Dnajb6-positive cells overlap only with a small portion of the HCN4-positive cells (Figure 2C and 2D). This is supported by a negative correlation between the expression level of Dnajb6 and Tbx3, as Tbx3 is one of the main transcriptional regulators of HCN4 expression in cardiac conduction system.^44, 45^ While these results may sound surprising, studies on isolated SAN cells reported dramatic variability in the density of HCN-formed “funny” current *I*f.^46-49^ In spontaneously beating cardiomyocytes isolated from the rabbit SAN, Monfredi et al. showed that *I*f density can range from 0 to ∼50 pA/pF and some the spontaneously beating SAN cells may have little to zero *I*f.^48^ The authors further observed SAN cells with lower *I*f current densities demonstrated a significantly greater sensitivity to inhibition of Ca^2+^ clock component of the SAN pacemaking machinery by cyclopiazonic acid, a moderate disruptor of Ca^2+^ cycling, in terms of beating rate slowing. The authors also noted that a relatively large cell population (21 of 90 cells) stopped beating when the sarcoplasmic reticulum pumping rate decreased in the presence of CPA, despite a relatively high *I*f density. Together with other studies,^33, 50^ these results may indicate a significant functional heterogeneity of pacemaker cells within the SAN in terms of their spontaneous beating rate, ion channel and calcium handling protein expression repertoire, and molecular mechanisms of their pacemaker activities. The latter was recently linked to the balance between the voltage and calcium components of the coupled-clock pacemaker system describing mechanisms of SAN automaticity.^51^ As summarized in details in our recent review,^52^ it was suggested that pacemaker cells, which primary rely on the Ca^2+^ clock, are more sensitive to the autonomic modulation through cAMP-mediated regulation of phosphorylation of Ca^2+^ handling proteins.^50^ This is in line with our findings indicating that Dnajb6 is mainly expressed in SAN cells with low HCN4 density (Figure 2D) and that Dnajb6 knock-out affects calcium homeostasis genes (Figure 6) and leads to abnormal autonomic regulation of the SAN (Figure 3E and Figure 4). Though our studies strongly support a crucial role of Dnajb6 in SAN automaticity and autonomic regulation of SAN pacemaking, detailed studies are needed to determine exact cellular and molecular pathways involved in these mechanisms.

### 3.3 A phenotype-based screening approach would facilitate the elucidation of molecular basis of SSS

Besides *dnajb6b*, our pilot forward genetic screen also suggested two additional candidate SSS genes like *cyth3a* and *vapal*, pending more experimental evidence to confirm their function. This forward genetic screening approach is scalable to the genome, which would generate a comprehensive list of candidate genes for SSS. Because there are at least 3 major cell types in the SAN region, including pacemaker cells in SAN that generate rhythm, paranodal areas and transition cells in the atrium that transmit the signal from pacemaker cells to govern coordinated contraction of the heart from atrium and then to the ventricle,^53, 54^ newly identified SSS genes could be categorized into different groups based on their expression pattern and phenotypic traits. We anticipate that systematic studies of these candidate genes identified from zebrafish will significantly advance our understanding of pathophysiology of SSS.

## 4. Materials and methods

### 4.1 Animals

All experiments were conducted in accordance with the National Institutes of Health Guide for the Care and Use of Laboratory Animals (NIH Pub. No. 80-23). All methods and protocols used in these studies have been approved by the Mayo Clinic Institutional Animal Care and Use Committee and by the Animal Care and Use Committee of University of Wisconsin-Madison, following the Guidelines for the Care and Use of Laboratory Animals published by the US National Institutes of Health (publication No. 85-23, revised 1996). The zebrafish (*Danio rerio*) WIK line was maintained under a 14-hour light/10-hour dark cycle at 28.5°C. All GBT lines were generated previously.^23-25^ The *Dnajb6* knockout (KO) mice, originally named *Dnajb6*^*tm1*.*1(KOMP)Vlcg*^, were generated from the Jackson Laboratory (Original catalog #018623). Briefly, the insertion of Velocigene cassette ZEN-Ub1 created a deletion sized 36,843 bp nucleotides spanning from the first to the last intron of the *Dnajb6* gene at the Chromosome 5 (Genome Build37) of the C57BL/6N mice. The mouse was subsequently bred to a ubiquitous Cre deletion mouse line for recombination of the LoxP sites that excised the neomycin selection cassette. The following genotyping PCR primers for the *Dnajb6* mutant mice were used: mutant primer F2, 5’-AAACTGCGCACTGTACCACC-3’ and mutant primer R2, 5’-CGGTCGCTACCATTACCAGT-3’ for detecting the mutant allele (predicted size of 700 bp); and WT primer F1, 5’-TACTCCAGCCCCACTCTTACTC-3’ and WT primer R1, 5’-ACTGCCCATCTTCTTCAACTTC-3’ for detecting the WT allele (predicted size of 300 bp).

### 4.2 Enrichment and cloning of 35 ZIC mutants

Zebrafish cardiac insertional (ZIC) mutants were identified and collected based on the mRFP expression in the embryonic heart from 2 to 4 days post-fertilization (dpf) and/or in the dissected adult heart at 6 months to 1 year of age. All ZIC lines, each with a single copy of GBT insertion, were obtained after 2 to 4 generations of outcrosses, guided by Southern blotting using the *GFP* probe primed to the GBT vector.^25^ A combination of 3 different methods including Inverse PCR, 5’-RACE and/or 3’-RACE were employed to clone the GBT transposon integration sites accordingly to previously published protocols.^23-25^

### 4.3 Zebrafish Electrocardiogram (ECG)

Microsurgery was operated under a dissection microscope to remove the silvery epithelial layer of the hypodermis one week before fish were subjected to the ECG.^29^ Fish were initially acclimated for one hour after transferred from the circulating fish facility to the laboratory bench, followed by anesthesia in the solution of pH 7.0 adjusted tricaine (MS-222, Sigma) at the concentration of 0.02% dissolved in E3 medium (containing 5 mM NaCl, 0.17 mM KCl, 0.33 mM CaCl2, and 0.33 mM MgSO4) for 6 minutes. Two minutes of ECG recording were then obtained with the ECG recording system, according to the instructions (ZS-200, iWorx Systems, Inc) and a recently published protocol.^29^ Initial ECG screens of ZIC heterozygous mutants were performed at 32°C using a temperature-controlled chamber set-up, made by covering the ECG recording system with a foam box. 6 to 25 fish per ZIC line were initially analyzed, depending on the fish availability. The ECG machine was held on top of a heating plate controlled by a heating machine. The subsequent ECG validation in the homozygous mutants was performed at room temperature (25°C). To analyze the ECG recording, ECG signals were amplified and filtered at 0.5 Hz high pass and 200 Hz low pass. ECG variables, including heart rate, P-wave amplitude, R-wave amplitude, and PP and RR intervals were calculated using an in-house Matlab code.^55^ A SA episode was defined in zebrafish when the PP interval is more than 1.5 seconds.

### 4.4 Mouse ECG and echocardiography

Mouse echocardiography and ECG measurements were performed according to a previously published protocol with modifications.^26, 27^ For ECG, mice were anesthetized with isoflurane (0.5% to 1.0% v/v) via a nose cone. Mice were placed on an ECG-heater board with 4 paws on individual electrodes. The ECG-heater board maintained the body temperature at 37°C. The ECG signal was amplified through an amplifier (Axon CNS digital 1440 A) and recorded using Chart 5 software. For each mouse, 10 min of ECG signal were recorded. Series of ECG parameters, including heart rate, P-wave amplitude, R-wave amplitude, PP/RR interval were calculated by an in-house Matlab code.^55^ For echocardiography, mice were anesthetized under light isoflurane (0.5% to 1.0% v/v) administered via a nose cone. Echocardiography gel was placed on the shaved chest, and the mouse heart was imaged with a 13-MHz probe using 2-dimensional echocardiography (GE Healthcare). All measurements were made by an independent operator to whom the study groups were masked.

### 4.5 Administration of autonomic response drugs

For zebrafish, 0.6 μg/g isoproterenol (Millipore Sigma, Cat# 1351005), 4 μg/g atropine (Millipore Sigma, Cat# A0132), and 0.3 μg/g carbachol (Millipore Sigma, cat# C4382) were administrated via intraperitoneal injection. For *in vivo* mouse studies, 0.2 mg/kg isoproterenol, 1 mg/kg atropine, and 0.3 mg/kg carbachol was injected intraperitoneally. For *ex vivo* mouse atrial studies, 100 nM isoproterenol, 2 μM atropine, and 300 nM carbachol was administered via superfusion for 10 - 20 min.

### 4.6 Antibody immunostaining

Heart samples harvested from mouse SAN tissues were embedded in a tissue freezing medium, followed by sectioning at 10 μm using a cryostat (Leica CM3050 S). The slides were subjected to immunostaining using a previously described protocol.^56^ The following antibodies were used: anti-HCN4 (Millipore, Cat#: AB5805) at 1:200, anti-Dnajb6 (Novus, Cat#: H00010049-M01) at 1:200, anti-Islet1 (abcam, Cat#: ab20670), anti-Tbx3 (abcam, Cat#: ab99302). All images were captured either using a Zeiss Axioplan II microscope equipped with ApoTome and AxioVision software (Carl Zeiss Microscopy) or a Zeiss LSM 780 confocal microscope.

### 4.7 Isolated mouse atrial preparations

The mouse atrial preparation was performed as previously described.^57^ After the mice were anesthetized with isoflurane, a mid-sternal incision was applied. The heart was then removed and cannulated to a custom made 21-gauge cannula. The heart was then perfused and superfused with oxygenated (95% O2, 5% CO2), 37°C modified Tyrode solution (in mM: 128.2 NaCl, 4.7 KCl, 1.19 NaH2PO4, 1.05 MgCl2, 1.3 CaCl2, 20.0 NaHCO3, and 11.1 glucose; pH=7.35±0.05). Lung, thymus and fat tissue was then removed. Perfusion was maintained under constant aortic pressure of 60-80 mmHg. After 10 min stabilization, the ventricles were dissected. The atrial were cut open as previously described.^58^ The medial limb of the crista terminalis was cut to open right atrial appendage. The preparation was superfused with Tyrode solution at a constant rate of ∼ 15 ml/min.

### 4.8 Optical mapping

High spatial (100×100 pixels, 60 ± 10μm per pixel) and temporal (1,000 – 3,000 frames/sec) resolution optical mapping of electrical activity was applied on the isolated mouse atrial preparations as previously described.^58, 59^ The isolated mouse atrial preparations were coronary and surface stained with voltage-sensitive dye RH-237 (1.25 mg/ml in dimethyl sulfoxide ThermoFisher Scientific, USA). Blebbistatin (10 μM, Tocris Bioscience, USA) was then applied to reduce the motion artifact. A 150-W halogen lamp (MHAB-150W, Moritex USA Inc., CA, USA) with band pass filter (530/40 nm) was used as excitation light source. The fluorescent light emitted from the preparation was recorded by a MiCAM Ultima-L camera (SciMedia, CA, USA) after a long-pass filter (>650 nm). The acquired fluorescent signal was digitized, amplified, and visualized using custom software (SciMedia, CA, USA). After 20-30 min stabilization, activation map was collected during baseline spontaneous rhythm. To estimate the pacemaker location and a possible pacemaker shift during autonomic stimulation, the originations of action potentials were plotted with orthogonal axes crossing at the inferior vena cava. The superior to inferior direction is along the ordinate. The lateral to media direction is along the abscissa. SAN recovery time (SANRT) was measured as the time-period between the last S1S1 pacing (10 Hz) beat and the first spontaneous beat. Corrected SANRT (SANRTc) was calculated as the difference between the SANRT and the resting cycle length measure before the SANRT pacing protocol. After baseline measurement, 100 nM isoproterenol was applied. Recordings were collected after 10 min which allows the stimulation to reach steady-state effect. Complete washout was then performed which is characterized by the recovery of the heart rate back to baseline values. Additional staining and blebbistatin was applied as needed. 300 nM carbachol then was applied. 2 μM atropine was used after protocols completed during carbachol stimulation.

### 4.9 RNA-seq data collection and analysis

Total RNA was extracted from the right atrium (RA) tissues of 1-year-old *Dnajb6*^*+/-*^ heterozygous mutant hearts and WT sibling controls. Six total samples (3 biological replicates for each genotype) were submitted for RNA sequencing (Azenta Life Science, NJ). Genes were considered to be differentially expressed between the two groups if they exhibited a greater than 2-fold change and an FDR of less than 0.05 according to the DESeq approach.^61^ Unsupervised hierarchical clustering was performed with Pearson correlation and scaled based on the fragments per kilobase of transcript per million mapped reads (FPKM) value using the pheatmap R package (https://github.com/raivokolde/pheatmap). The gene lists of interest were annotated by IPA (QIAGEN) (http://www.ingenuity.com/). We queried the IPA with the gene list of interest to map and generate putative biological processes/functions, networks, and pathways based on the manually curated knowledge database of molecular interactions extracted from the public literature. The enriched pathways and gene networks were generated using both direct and indirect relationships/connectivity. These pathways and networks were ranked by their enrichment score, which measures the probability that the genes were included in a network by chance.

### 4.10 Quantitative reverse transcription (RT) PCR

Total RNA was extracted from ∼2 mg of right atrium (RA) tissues of 1-year-old *Dnajb6*^*+/-*^ heterozygous mutant hearts and WT sibling controls using Trizol reagent (ThermoFisher Scientific) following the manufacturer’s instruction. Approximately 1 μg total RNA was used for reverse transcription (RT) and cDNA synthesis using Superscript III First-Strand Synthesis System (ThermoFisher Scientific). Real-time quantitative RT-PCR was run in 96-well optical plates (ThermoFisher Scientific) using an Applied Biosystem VAii 7 System (ThermoFisher Scientific). Gene expression levels were normalized using the expression level of glyceraldehyde 3-phosphate dehydrogenase (*gapdh)* by –ΔΔCt (cycle threshold) values. All quantitative RT-PCR primer sequences were listed in Supplemental Table 2.

### 4.10 Statistics

No sample sizes were calculated before performing the experiments. No animals were excluded for analysis. Unpaired 2-tailed student’s *t*-test was used to compare 2 groups. One-way Analysis of Variance (ANOVA) or Kruskal-Wallis test followed by post hoc Tukey’s test was used for comparing 3 and more groups. Chi-square test was used for rate comparison. *P* values less than 0.05 was considered statistically significant. For dot plot graphs, values are displayed as mean ± standard deviation (SD). Sample size (N) represents animal number, otherwise specifically designated as biological or technical replicates. All statistical analyses were conducted with the Graphpad Prism 7 and/or R Statistical Software Version 3.6.1.

## Acknowledgments

We thank Beninio Gores and Kashia Stragey for managing zebrafish facility and Ronald H. May for murine echocardiography.

## Sources of Funding

This work was supported in part by grants from the Mayo Foundation to X.X., Grants from YG Li (Science and Technology Innovation Action Plan of Shanghai, experimental animal research project 201409005600), NIH R01HL141214, American Heart Association 16SDG29120011, and the Wisconsin Partnership Program 4140 to A.V.G., and American Heart Association 17POST33370089 and American Heart Association Career Development Award 846898 to D. L.

## Notes

### Competing Interest Statement

The authors have declared no competing interest.

