## Supplemental Figures and Tables for "A phenotype-based forward genetic screen identifies *Dnajb6* as a sick sinus syndrome gene"

### Slide 1
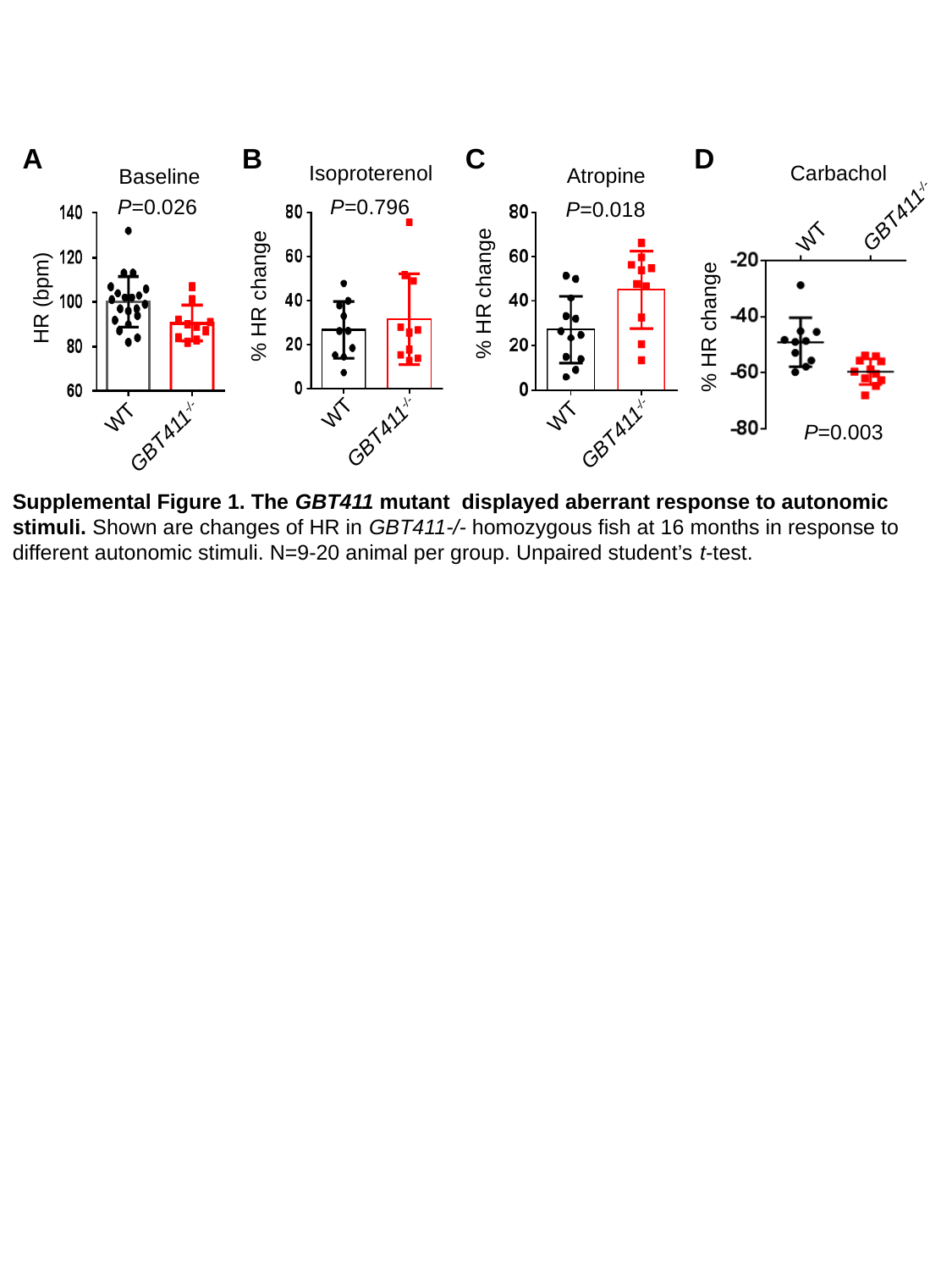

A
B
C
D
Carbachol
Isoproterenol
Atropine
Baseline
P=0.026
P=0.796
P=0.018
GBT411^-/-^
WT
% HR change
% HR change
HR (bpm)
% HR change
WT
WT
WT
GBT411^-/-^
P=0.003
GBT411^-/-^
GBT411^-/-^
Supplemental Figure 1. The GBT411 mutant displayed aberrant response to autonomic stimuli. Shown are changes of HR in GBT411-/- homozygous fish at 16 months in response to different autonomic stimuli. N=9-20 animal per group. Unpaired student’s t-test.

### Slide 2
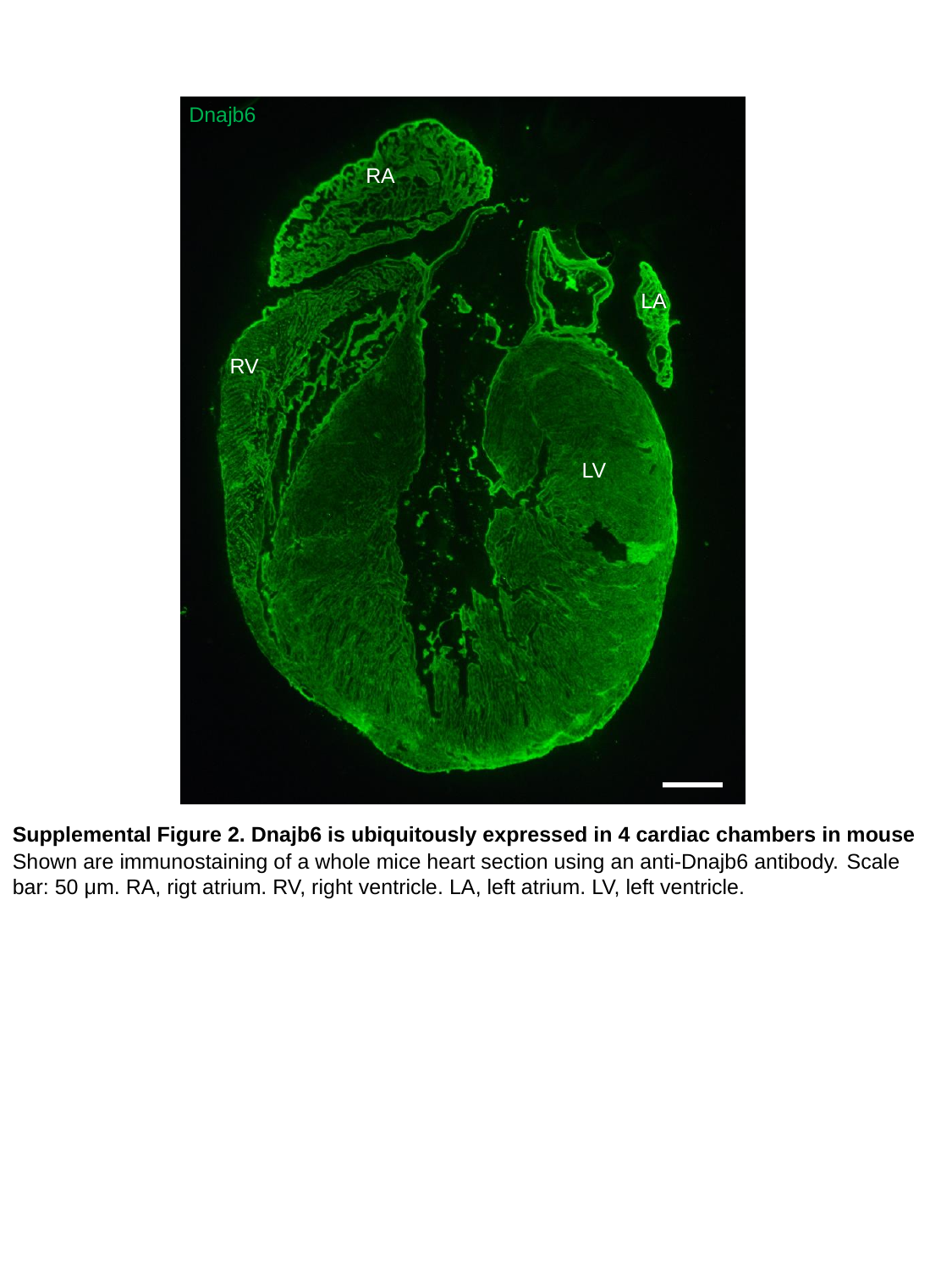

Dnajb6
RA
LA
RV
LV
Supplemental Figure 2. Dnajb6 is ubiquitously expressed in 4 cardiac chambers in mouse
Shown are immunostaining of a whole mice heart section using an anti-Dnajb6 antibody. Scale bar: 50 μm. RA, rigt atrium. RV, right ventricle. LA, left atrium. LV, left ventricle.

### Slide 3
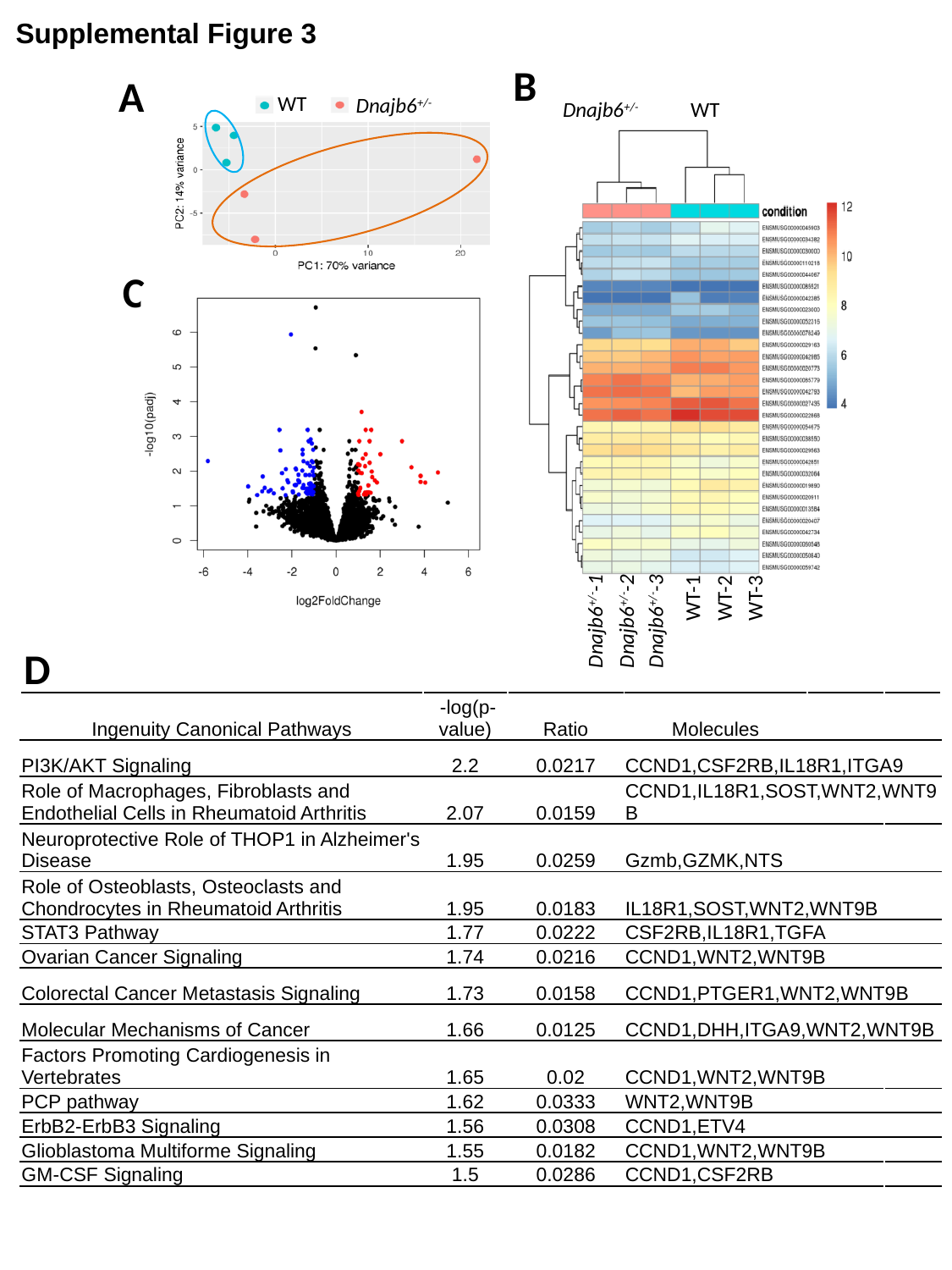

Supplemental Figure 3
B
A
WT
 Dnajb6^+/-^
 Dnajb6^+/-^
WT
C
WT-2
WT-1
WT-3
Dnajb6^+/-^-1
Dnajb6^+/-^-3
Dnajb6^+/-^-2
D
| Ingenuity Canonical Pathways | -log(p-value) | Ratio | Molecules |
| --- | --- | --- | --- |
| PI3K/AKT Signaling | 2.2 | 0.0217 | CCND1,CSF2RB,IL18R1,ITGA9 |
| Role of Macrophages, Fibroblasts and Endothelial Cells in Rheumatoid Arthritis | 2.07 | 0.0159 | CCND1,IL18R1,SOST,WNT2,WNT9B |
| Neuroprotective Role of THOP1 in Alzheimer's Disease | 1.95 | 0.0259 | Gzmb,GZMK,NTS |
| Role of Osteoblasts, Osteoclasts and Chondrocytes in Rheumatoid Arthritis | 1.95 | 0.0183 | IL18R1,SOST,WNT2,WNT9B |
| STAT3 Pathway | 1.77 | 0.0222 | CSF2RB,IL18R1,TGFA |
| Ovarian Cancer Signaling | 1.74 | 0.0216 | CCND1,WNT2,WNT9B |
| Colorectal Cancer Metastasis Signaling | 1.73 | 0.0158 | CCND1,PTGER1,WNT2,WNT9B |
| Molecular Mechanisms of Cancer | 1.66 | 0.0125 | CCND1,DHH,ITGA9,WNT2,WNT9B |
| Factors Promoting Cardiogenesis in Vertebrates | 1.65 | 0.02 | CCND1,WNT2,WNT9B |
| PCP pathway | 1.62 | 0.0333 | WNT2,WNT9B |
| ErbB2-ErbB3 Signaling | 1.56 | 0.0308 | CCND1,ETV4 |
| Glioblastoma Multiforme Signaling | 1.55 | 0.0182 | CCND1,WNT2,WNT9B |
| GM-CSF Signaling | 1.5 | 0.0286 | CCND1,CSF2RB |

### Slide 4
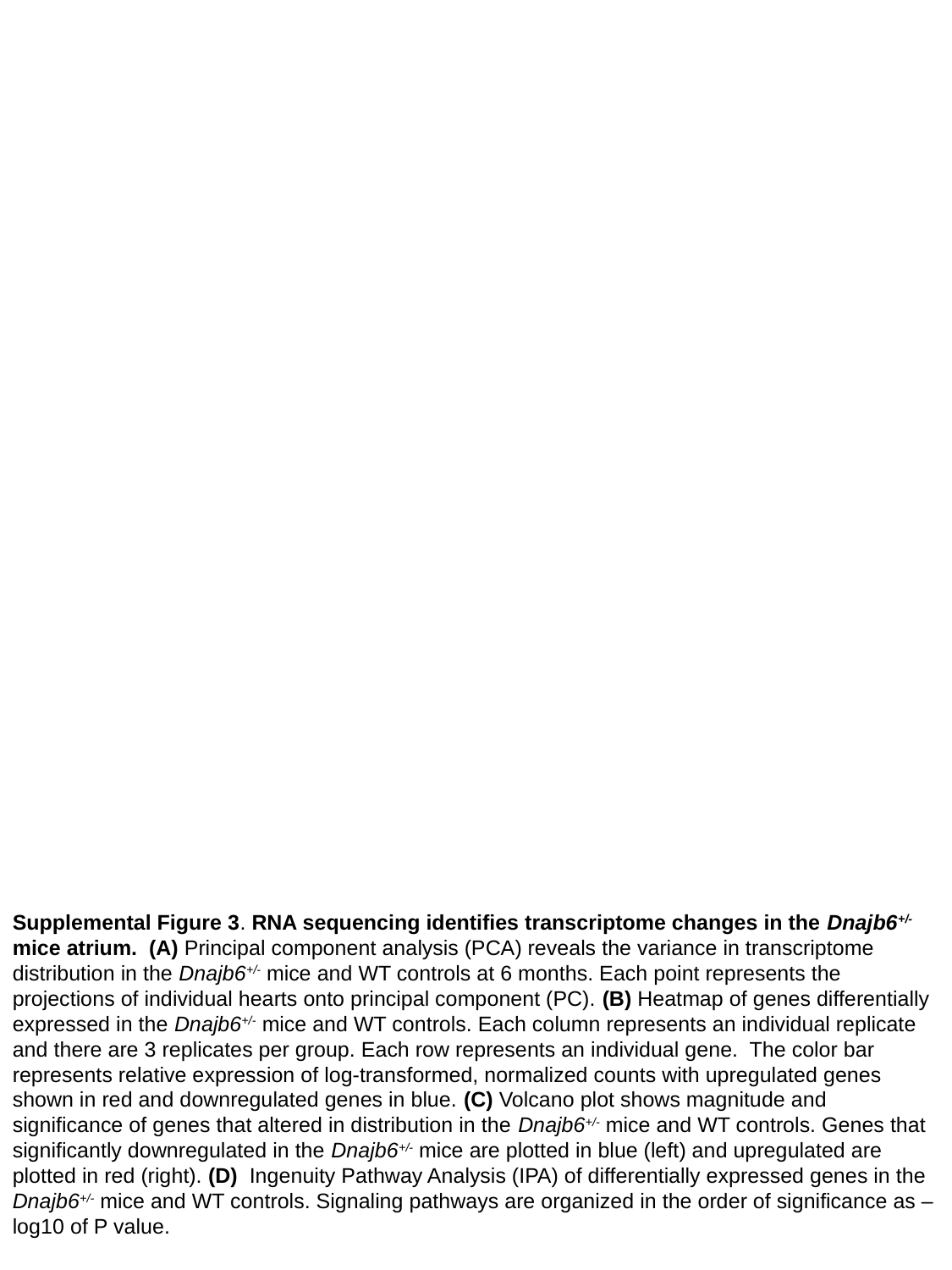

Supplemental Figure 3. RNA sequencing identifies transcriptome changes in the Dnajb6^+/-^ mice atrium. (A) Principal component analysis (PCA) reveals the variance in transcriptome distribution in the Dnajb6^+/-^ mice and WT controls at 6 months. Each point represents the projections of individual hearts onto principal component (PC). (B) Heatmap of genes differentially expressed in the Dnajb6^+/-^ mice and WT controls. Each column represents an individual replicate and there are 3 replicates per group. Each row represents an individual gene. The color bar represents relative expression of log-transformed, normalized counts with upregulated genes shown in red and downregulated genes in blue. (C) Volcano plot shows magnitude and significance of genes that altered in distribution in the Dnajb6^+/-^ mice and WT controls. Genes that significantly downregulated in the Dnajb6^+/-^ mice are plotted in blue (left) and upregulated are plotted in red (right). (D)  Ingenuity Pathway Analysis (IPA) of differentially expressed genes in the Dnajb6^+/-^ mice and WT controls. Signaling pathways are organized in the order of significance as –log10 of P value.

### Slide 5
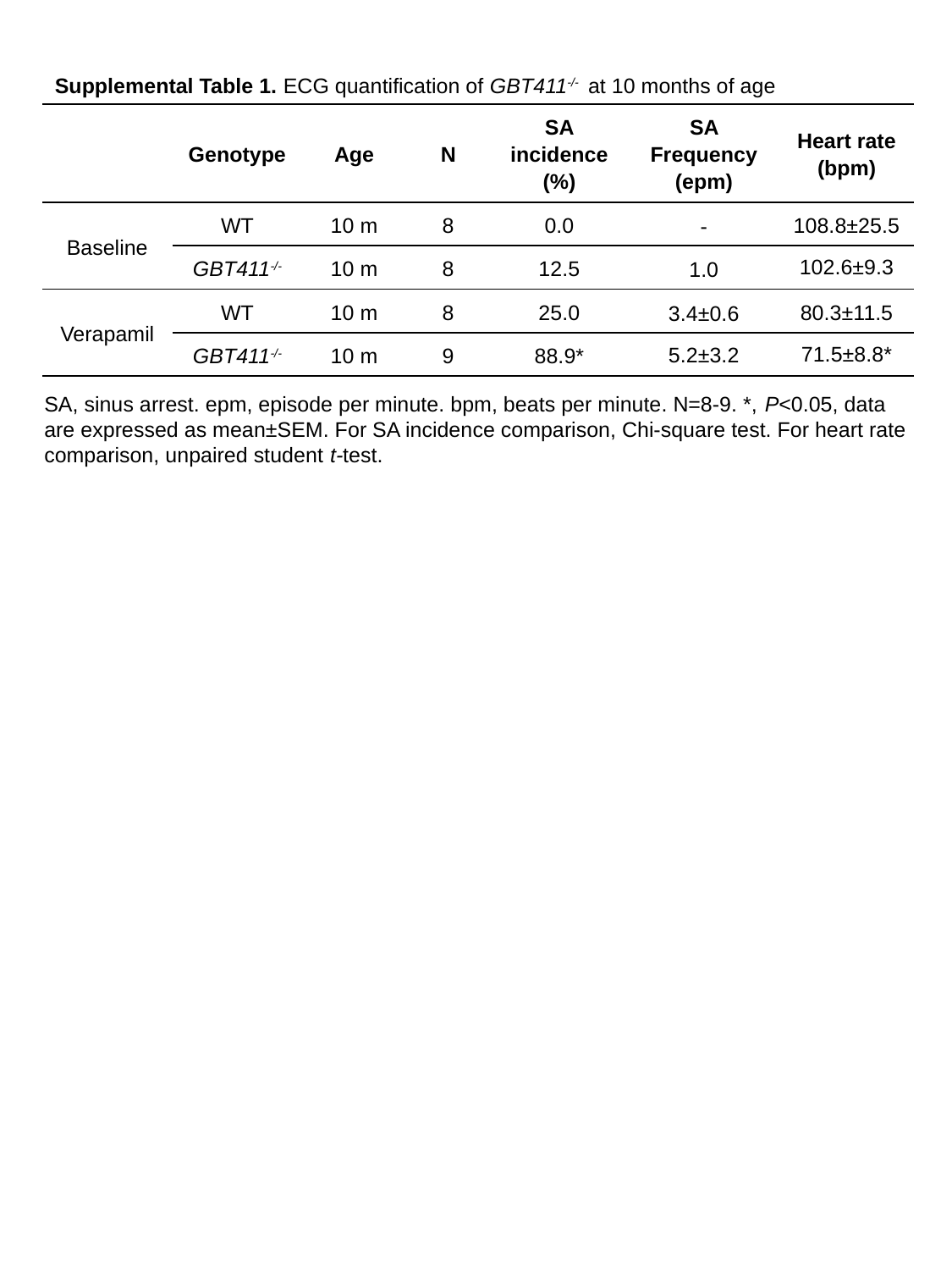

Supplemental Table 1. ECG quantification of GBT411^-/-^ at 10 months of age
| | Genotype | Age | N | SA incidence (%) | SA Frequency (epm) | Heart rate (bpm) |
| --- | --- | --- | --- | --- | --- | --- |
| Baseline | WT | 10 m | 8 | 0.0 | - | 108.8±25.5 |
| | GBT411^-/-^ | 10 m | 8 | 12.5 | 1.0 | 102.6±9.3 |
| Verapamil | WT | 10 m | 8 | 25.0 | 3.4±0.6 | 80.3±11.5 |
| | GBT411^-/-^ | 10 m | 9 | 88.9\* | 5.2±3.2 | 71.5±8.8\* |
SA, sinus arrest. epm, episode per minute. bpm, beats per minute. N=8-9. *, P<0.05, data are expressed as mean±SEM. For SA incidence comparison, Chi-square test. For heart rate comparison, unpaired student t-test.

### Slide 6
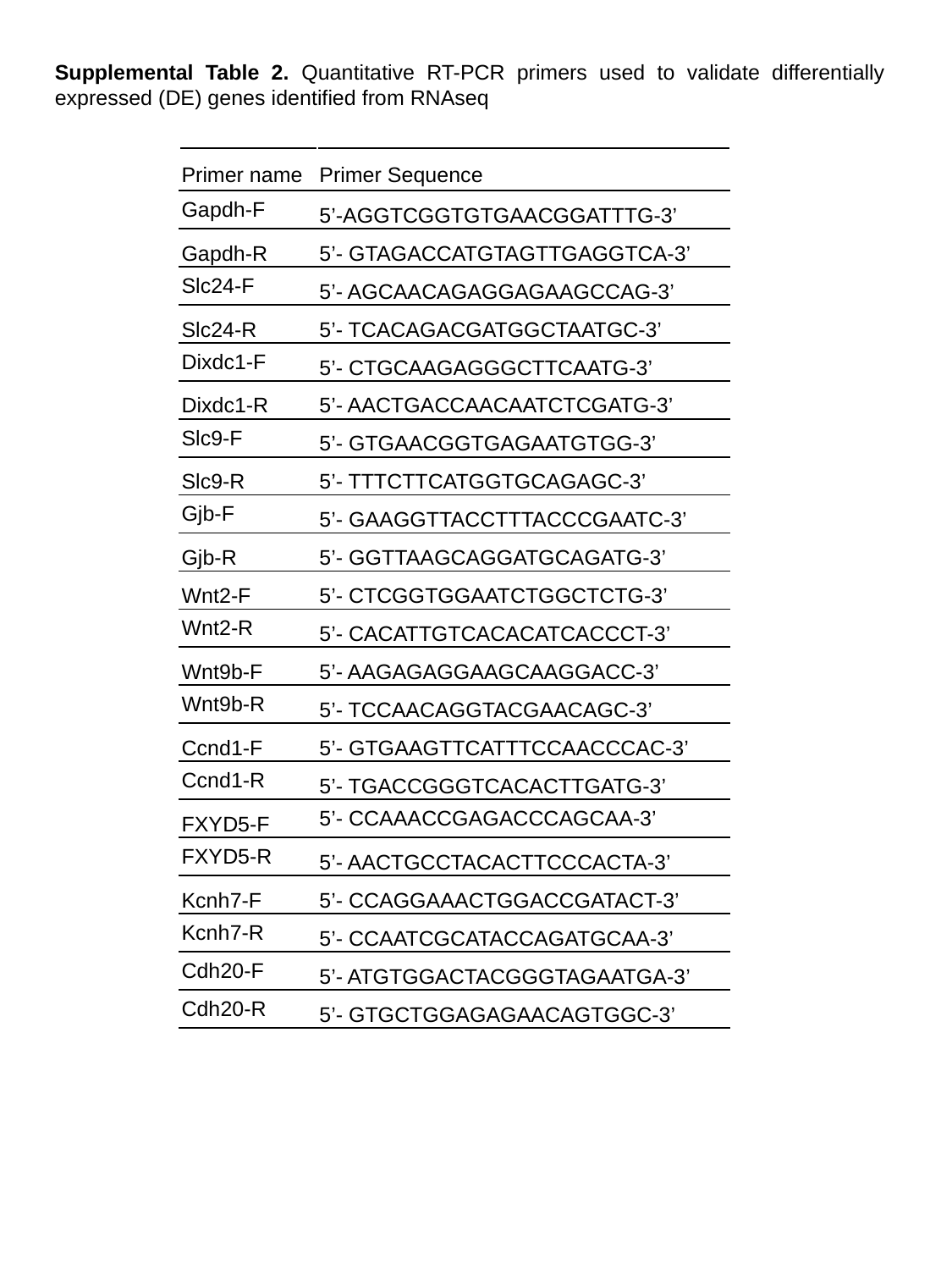

Supplemental Table 2. Quantitative RT-PCR primers used to validate differentially expressed (DE) genes identified from RNAseq
| Primer name | Primer Sequence |
| --- | --- |
| Gapdh-F | 5’-AGGTCGGTGTGAACGGATTTG-3’ |
| Gapdh-R | 5’- GTAGACCATGTAGTTGAGGTCA-3’ |
| Slc24-F | 5’- AGCAACAGAGGAGAAGCCAG-3’ |
| Slc24-R | 5’- TCACAGACGATGGCTAATGC-3’ |
| Dixdc1-F | 5’- CTGCAAGAGGGCTTCAATG-3’ |
| Dixdc1-R | 5’- AACTGACCAACAATCTCGATG-3’ |
| Slc9-F | 5’- GTGAACGGTGAGAATGTGG-3’ |
| Slc9-R | 5’- TTTCTTCATGGTGCAGAGC-3’ |
| Gjb-F | 5’- GAAGGTTACCTTTACCCGAATC-3’ |
| Gjb-R | 5’- GGTTAAGCAGGATGCAGATG-3’ |
| Wnt2-F | 5’- CTCGGTGGAATCTGGCTCTG-3’ |
| Wnt2-R | 5’- CACATTGTCACACATCACCCT-3’ |
| Wnt9b-F | 5’- AAGAGAGGAAGCAAGGACC-3’ |
| Wnt9b-R | 5’- TCCAACAGGTACGAACAGC-3’ |
| Ccnd1-F | 5’- GTGAAGTTCATTTCCAACCCAC-3’ |
| Ccnd1-R | 5’- TGACCGGGTCACACTTGATG-3’ |
| FXYD5-F | 5’- CCAAACCGAGACCCAGCAA-3’ |
| FXYD5-R | 5’- AACTGCCTACACTTCCCACTA-3’ |
| Kcnh7-F | 5’- CCAGGAAACTGGACCGATACT-3’ |
| Kcnh7-R | 5’- CCAATCGCATACCAGATGCAA-3’ |
| Cdh20-F | 5’- ATGTGGACTACGGGTAGAATGA-3’ |
| Cdh20-R | 5’- GTGCTGGAGAGAACAGTGGC-3’ |
